# Multiplexed identification of RAS paralog imbalance as a driver of lung cancer growth

**DOI:** 10.1101/2021.07.08.451571

**Authors:** Rui Tang, Emily G. Shuldiner, Marcus Kelly, Christopher W. Murray, Jess D. Hebert, Laura Andrejka, Min K. Tsai, Nicholas W. Hughes, Mitchell I Parker, Hongchen Cai, Yao-Cheng Li, Geoffrey M. Wahl, Roland L. Dunbrack, Peter K. Jackson, Dmitri A. Petrov, Monte M. Winslow

**Affiliations:** Department of Genetics, Stanford University School of Medicine, Stanford, CA, USA; Department of Biology, Stanford University, Stanford, CA, USA; Cancer Biology Program, Stanford University School of Medicine, Stanford, CA, USA; Baxter Laboratories, Stanford University School of Medicine, Stanford, CA, USA; Molecular Therapeutics Program, Institute for Cancer Research, Fox Chase Cancer Center, Philadelphia, PA, USA; Gene Expression Laboratory, The Salk Institute for Biological Studies, La Jolla, CA, USA; Department of Pathology, Stanford University School of Medicine, Stanford, CA, USA

**Author notes:** These authors contributed equally (Rui Tang, Emily G. Shuldiner). Corresponding author: Monte M. Winslow.

## Abstract

Oncogenic *KRAS* mutations occur in approximately 30% of lung adenocarcinoma. Despite several decades of effort, oncogenic KRAS-driven lung cancer remains difficult to treat, and our understanding of the positive and negative regulators of RAS signaling is incomplete. To uncover the functional impact of diverse KRAS-interacting proteins on lung cancer growth *in vivo*, we used multiplexed somatic CRISPR/Cas9-based genome editing in genetically engineered mouse models with tumor barcoding and high-throughput barcode sequencing. Through a series of CRISPR/Cas9 screens in autochthonous lung tumors, we identified HRAS and NRAS as key suppressors of KRAS^G12D^-driven tumor growth *in vivo* and confirmed these effects in oncogenic KRAS-driven human lung cancer cell lines. Mechanistically, RAS paralogs interact with oncogenic KRAS, suppress KRAS-KRAS interactions, and reduce downstream ERK signaling. HRAS mutations identified in KRAS-driven human tumors partially abolished this effect. Comparison of the tumor-suppressive effects of HRAS and NRAS in KRAS- and BRAF-driven lung cancer models confirmed that RAS paralogs are specific suppressors of oncogenic KRAS-driven lung cancer *in vivo*. Our study outlines a technological avenue to uncover positive and negative regulators of oncogenic KRAS-driven cancer in a multiplexed manner *in vivo* and highlights the role of RAS paralog imbalance in oncogenic KRAS-driven lung cancer.

## INTRODUCTION

The RAS family genes *KRAS*, *HRAS* and *NRAS* are frequently mutated across cancers, and *KRAS* mutations occur in approximately 30% of lung adenocarcinomas^1–3^. RAS proteins are small GTPases that switch between a GTP-bound active state and GDP-bound inactive state in response to upstream growth signaling^4^. RAS proteins regulate multiple downstream signaling pathways which control proliferation. Hotspot oncogenic mutations in codons 12, 13, and 61 reduce GTP hydrolysis and increase the fraction of RAS proteins in the GTP-bound state, which results in constitutive activation and widespread changes in RAS protein-protein interactions^5, 6^. These changes result in hyper-activation of RAS effector pathways, driving cellular transformation and tumorigenesis^7, 8^. Oncogenic KRAS therefore represents a key node in growth factor-induced signaling and a critical target for therapeutic intervention in lung adenocarcinoma. However, despite tremendous effort, the development of targeted therapies for oncogenic KRAS-driven tumors has proven challenging^9^.

Genetic and proteomic mapping has revealed that KRAS interacts with a large network of proteins^10, 11^. These KRAS-interacting proteins include canonical regulators and effectors, as well as many proteins that remain poorly understood in the context of oncogenic KRAS-driven lung cancer. Much of our understanding of RAS signaling has stemmed from diverse cellular and cell-free systems^12–14^. Thus, while recent studies have mapped KRAS protein-protein interaction networks and identified synthetic lethal interactions with oncogenic KRAS in human cell lines^10, 11, 15, 16^, it remains difficult to assess the relevance of these biochemical and genetic interactions to cancer growth *in vivo*. Genetically engineered mouse models of oncogenic KRAS-driven cancer uniquely recapitulate autochthonous tumor growth and have contributed to our understanding of KRAS signaling^17^. However, the development and use of such models has traditionally been insufficiently scalable to broadly assess modifiers of KRAS-driven tumor growth. The ability to uncover functional components of RAS signaling that affect lung cancer growth *in vivo* in a multiplexed manner would accelerate our understanding of RAS biology and could aid in the development of pharmacological strategies to counteract hyperactivated KRAS.

To enable the analysis of genetic modifiers of lung tumor growth *in vivo*, we recently integrated somatic CRISPR/Cas9-based genome editing with tumor barcoding and high-throughput barcode sequencing (Tuba-seq)^18–20^. This approach allows precise quantification of the effect of inactivating panels of genes of interest on lung tumor initiation and growth in a multiplexed manner. By employing Tuba-seq to assess the functions of KRAS-interacting proteins nominated by unbiased affinity purification/mass spectrometry (AP/MS), we show that wild-type HRAS and NRAS suppress the growth of oncogenic KRAS-driven lung adenocarcinoma. Competition between oncogenic KRAS and wild-type HRAS diminishes KRAS-KRAS interaction and suppresses downstream signaling. *In vivo* screening across multiple oncogenic contexts revealed that HRAS and NRAS specifically suppress the growth of tumors driven by oncogenic KRAS. Our study reveals that RAS paralog imbalance is a driver of oncogenic KRAS-driven lung cancer.

## RESULTS

### Selection of candidate KRAS-interacting proteins to assess *in vivo*

To identify putative KRAS-interacting proteins that could affect oncogenic KRAS-driven lung tumor growth *in vivo*, we integrated previous proteomic data from AP/MS studies with gene expression data from cancer cells from autochthonous mouse models (**Figure 1a**)^10, 21^. We prioritized a list of candidate genes according to the probability of their protein products interacting with KRAS, their mRNA expression in mouse models of oncogenic KRAS^G12D^-driven lung cancer, and the probability of their protein products interacting with other RAS GTPases (**Figure 1b-c, Figure S1a-d**)^10, 21^. We selected 13 proteins that represent diverse aspects of RAS biology, including RAS paralogs (HRAS, NRAS – which were supported by the identification of paralog-specific peptides), RAS regulators (RASGRF2, RAP1GDS1)^22, 23^, a RAS farnesyltransferase (FNTA)^24, 25^, and RAS effectors (RAF1, RGL2)^26, 27^, as well as several other proteins whose functions in RAS signaling are understudied. Analysis of human lung adenocarcinoma genomic data showed that while most of these candidate genes trend to be more often amplified in human adenocarcinoma, *NRAS*, *HRAS*, and *ALDH1A1* also have deep genomic deletions (**Figure S1e**)^28^. Interestingly, some of these proteins bound preferentially to either GTP- or GDP-bound KRAS, while others seemed to interact with KRAS independent of its nucleotide state **(Figure 1c)**.

**Figure 1.**
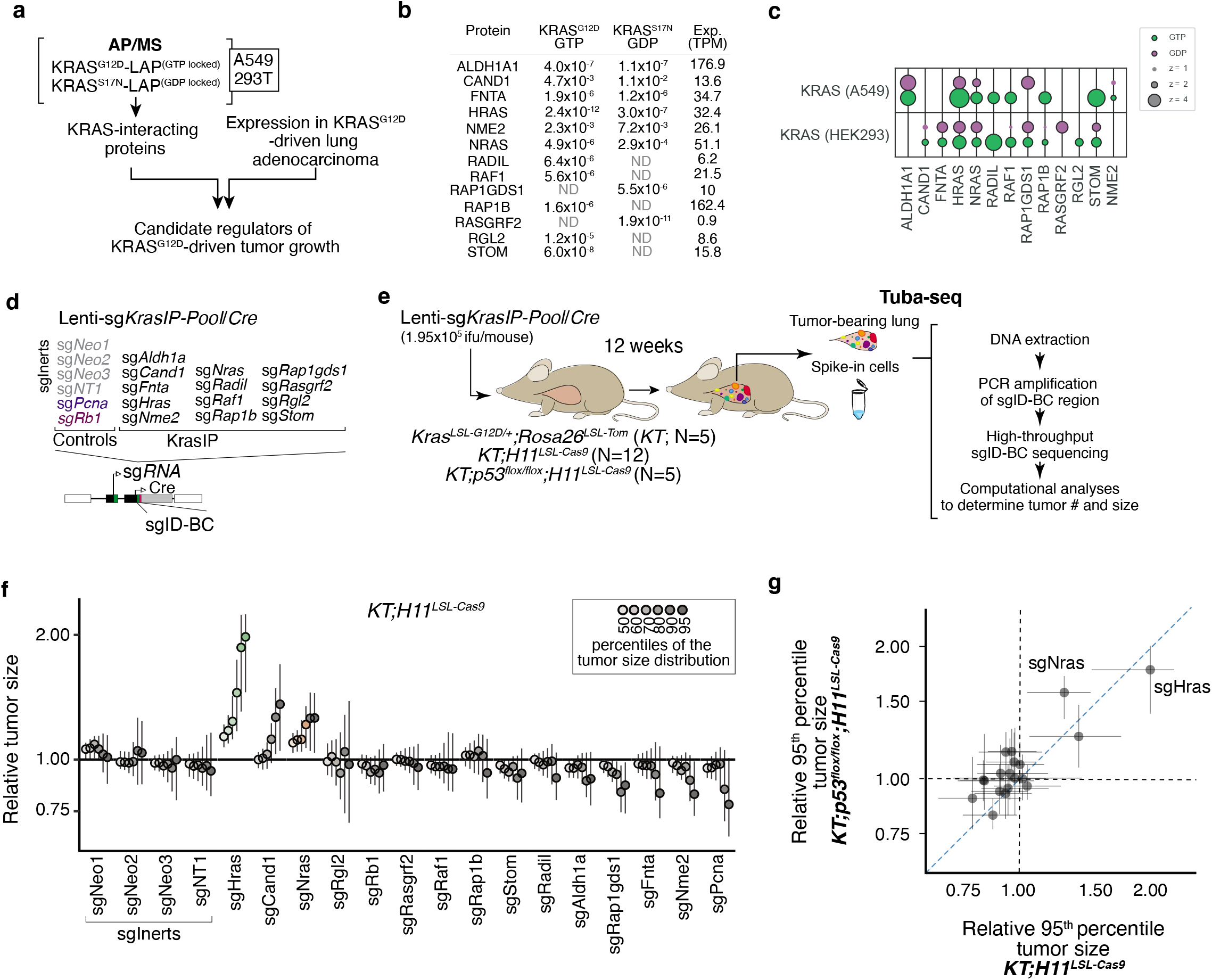
Multiplexed identification of KRAS-interacting proteins that impact KRAS^G12D^-driven lung cancer growth in vivo. **a.** Candidate mediators of KRAS-driven lung tumor growth were identified on the basis of their interactions with GTP- and GDP-locked Kras in multiple AP/MS-based protein-protein interaction screens and their expression in a mouse model of Kras-driven lung adenocarcinoma. **b.** Selected KRAS-interacting proteins interact with either GTP- or GDP-locked KRAS (shown as NSAF in A549 cells) and their homolog is expressed in KRAS^G12D^-driven lung cancer (shown as TPM). **c.** Bubble plot of two AP/MS experiments with GTP- and GDP-locked mutant GTPases as baits (rows), showing the enrichment of selected candidate KRAS-interacting proteins (columns). Dark borders indicate FDR < 0.05. **d.** Schematic of tumor initiation with a pool of barcoded Lenti-sgRNA/Cre vectors (Lenti-sgKrasIP-Pool/Cre). The lentiviral pool includes four Inert sgRNAs that are either non-targeting (NT) or target a functionally inert locus (Neo1-3, targeting *NeoR* in the *R26^LSL-tdTomato^* allele). Each barcoded lentiviral vector contains an sgRNA, Cre, and a two-component barcode composed of an sgRNA identifier (sgID) and a random barcode (BC). This design allows inactivation of multiple target genes in parallel followed by quantification of the resulting tumor size distributions through high-throughput sgID-BC sequencing. **e.** Tumors were initiated in cohorts of *KT*, *KT;H11^LSL-Cas9^* and *KT;p53^flox/flox^;H11^LSL-Cas9^* mice through intratracheal delivery of Lenti-sgKrasIP-Pool/Cre. Tuba-seq was performed on each tumor-bearing lung 12 weeks after initiation, followed by analyses of sgID-BC sequencing data to characterize the effects of inactivating each gene. **f.** Tumor sizes at indicated percentiles for each sgRNA relative to the size of sgInert-containing tumors at the corresponding percentiles in *KT;H11^LSL-Cas9^* mice. Genes are ordered by 95^th^ percentile tumor size, with sgInerts on the left. sgInerts are in gray, and the line at y=1 indicates no effect relative to sgInert. Error bars indicate 95% confidence intervals. Percentiles that are significantly different from sgInert (two-sided FDR-adjusted p < 0.05) are in color. Confidence intervals and P-values were calculated by bootstrap resampling. **g.** Comparison of 95^th^ percentile tumor size for each sgRNA relative to the size the 95^th^ percentile tumor size of sgInert-containing tumors in *KT;H11^LSL-Cas9^* mice versus *KT;p53^flox/flox^;H11^LSL-Cas9^* mice. Error bars indicate 95% confidence intervals calculated by bootstrap resampling.

### Identification of KRAS-interacting proteins that impact lung tumor growth *in vivo*

Given that KRAS-interacting proteins could have either positive or negative effects on signaling and tumor growth, we first assessed whether Tuba-seq is capable of detecting gene-targeting events that have deleterious effects on tumor fitness. We initiated tumors in *Kras^LSL-G12D/+^;Rosa26^LSL-tdTomato^;H11^LSL-Cas^*^9^ (*KT;H11^LSL-Cas9^*) and control *KT* mice with a pool of barcoded Lenti-sgRNA/Cre vectors encoding sgRNAs targeting two essential genes (*Pcna* and *Rps19*), a known tumor suppressor (*Apc*)^20, 29^, and several inert sgRNAs (Lenti-sgEssential/Cre; **Figure S2a**). After 12 weeks of tumor growth, we performed Tuba-seq on bulk tumor-bearing lungs and quantified the number and size of tumors initiated with each Lenti-sgRNA/Cre vector (**Figure S2b**). By incorporating measures of tumor number and size, we could confidently identify genetic deficiencies that reduced tumor fitness (**Figure S2c-g and Methods**).

To quantify the impact of inactivating our panel of KRAS-interacting proteins on oncogenic KRAS^G12D^-driven lung tumor growth *in vivo*, we generated a pool of barcoded Lenti-sgRNA/Cre vectors targeting the genes that encode these proteins, as well as sgInert control vectors (Lenti-sg*KrasIP/Cre*; **Figure 1d**). Given the importance of farnesylation in KRAS localization and signaling, sgRNA targeting *Fnta* served as a control for KRAS dependency^30, 31^. We initiated tumors with the Lenti-sg*KrasIP/Cre* pool in *KT;H11^LSL-Cas9^* and *KT* mice and calculated metrics of tumor size and number after 12 weeks of tumor growth (**Figure 1e**). To our surprise, inactivation of the *Kras* paralogs *Hras* and *Nras* had the most dramatic effect on tumor growth. Inactivation of *Cand1* also increased tumor size, while deletion of several genes including *Fnta*, *Nme2*, *Rap1gds1*, and *Aldh1a* decreased tumor size and/or number, suggesting reduced cancer cell fitness (**Figure 1f** and **S3a-d**).

Given the fundamental importance of the p53 tumor suppressor in oncogenic KRAS-driven lung cancer, as well as previous data suggesting crosstalk between RAS and p53 signaling^19, 32, 33^, we determined whether p53 deficiency changed the impact of inactivating KRAS-interacting proteins on tumor growth. We initiated tumors with the Lenti-sg*KrasIP/Cre* pool in *Kras^LSL-G12D/+^;Rosa26^LSL-tdTom^;Trp53^flox/flox^;H11^LSL-tdTom^ (KT;Trp53^flox/flox^;H11^LSL-Cas^*^9^) mice and performed Tuba-seq after 12 weeks of tumor growth (**Figure 1e**). The effects of inactivating each gene encoding a KRAS-interacting protein on tumor size, tumor number, and overall tumor burden were generally consistent between the p53-proficient and -deficient settings (**Figure 1g, Figure S3e-h**). Notably, the inactivation of either *Hras* or *Nras* also significantly increased growth of p53-deficient tumors (**Figure 1g, Figure S3e)**. Collectively, these results suggest that HRAS and NRAS are tumor suppressors within *in vivo* models of oncogenic KRAS-driven lung cancer, while several other KRAS-interacting proteins, including CAND1, ALDH1A, and NME2, have less consistent effects on tumor growth between p53-proficient and -deficient backgrounds (**Figure S3e-h**).

### Validation of HRAS and NRAS as suppressors of oncogenic KRAS-driven lung tumor growth

To further validate the effect of inactivating six top candidate genes (*Hras, Nras, Cand1, Aldh1a, Fnta, and Nme2*) on oncogenic KRAS-driven tumor growth *in vivo* and confirm that these results are driven by on-target effects, we generated and barcoded three Lenti-sgRNA/Cre vectors targeting each gene. To contextualize the effect of *Hras* and *Nras* inactivation on lung tumor growth relative to established tumor suppressors we included vectors targeting three established tumor suppressors (*Lkb1, Rbm10*, and *Rb1*) in this pool (Lenti-sgValidation/Cre; **Figure 2a**)^18, 20, 34^. We initiated tumors with the Lenti-sgValidation/Cre pool in *KT;H11^LSL-Cas9^* and *KT* mice and assessed metrics of tumor initiation and growth 12 weeks after tumor initiation (**Figure 2b-c**). Targeting *Fnta* with all three sgRNAs consistently reduced growth fitness, while the impact of inactivating *Aldh1a* and *Nme2* was more variable (**Figure 2d, Figure S4**). Most importantly, all three sgRNAs targeting *Hras* and all three sgRNAs targeting *Nras* significantly increased tumor growth (**Figure 2d-e, Figure S4b**). Notably, *Hras* inactivation increase tumor growth to a similar extent as inactivation of the *Rb1* and *Rbm10* tumor suppressors (**Figure 2d, Figure S4b**). These results suggest a potentially pivotal role for wild-type HRAS and NRAS in constraining oncogenic KRAS-driven lung tumor growth *in vivo*.

**Figure 2.**
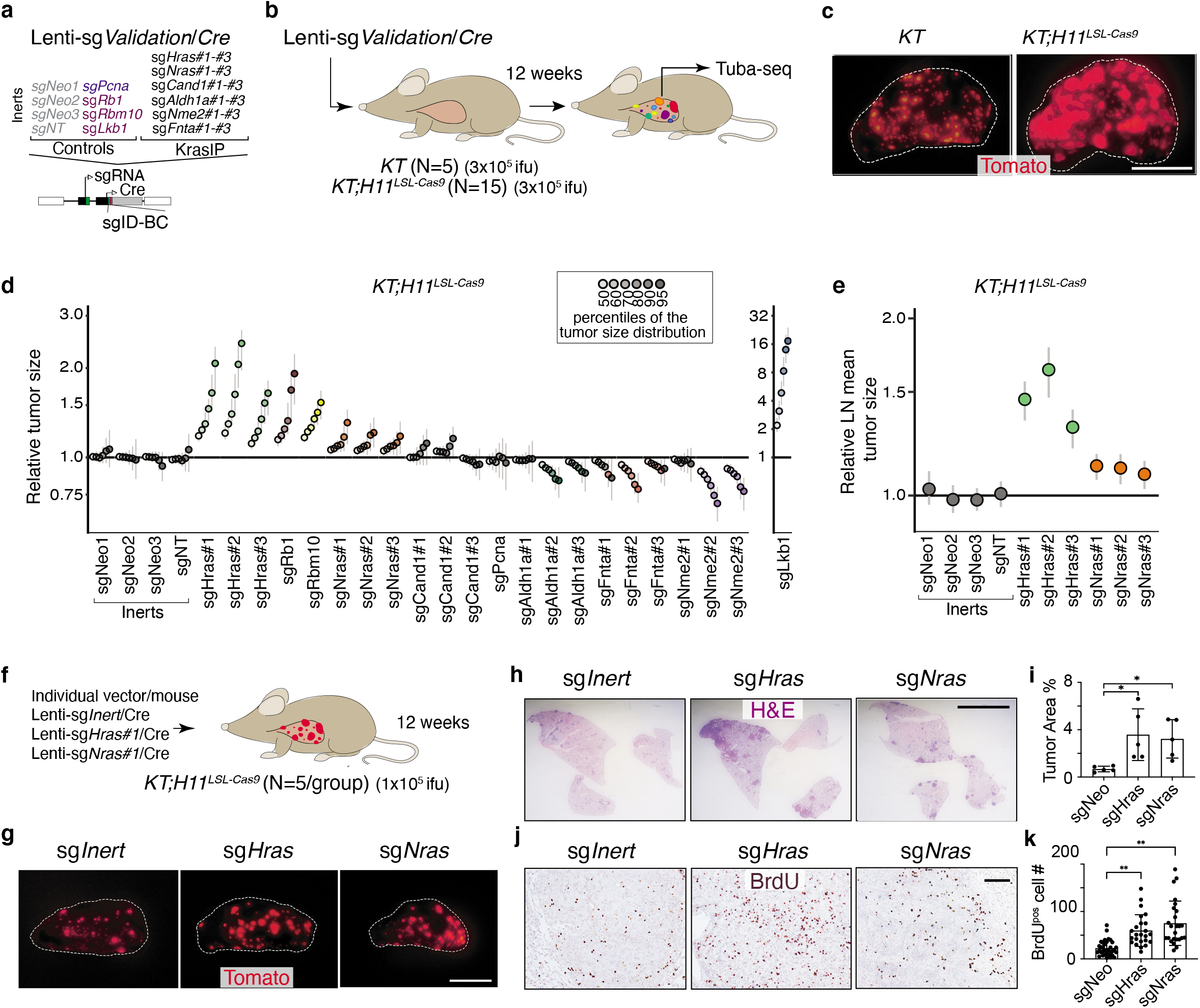
HRAS and NRAS are potent suppressors of KRAS^G12D^-driven lung cancer growth in vivo. **a,b.** A pool of barcoded Lenti-sgRNA/Cre vectors (Lenti-sgValidation/Cre) targeting candidate mediators of KRAS-driven lung tumor growth identified in the initial KRAS-interacting protein Tuba-seq screen was used to initiate tumors in validation cohorts of *KT* and *KT;H11^LSL-Cas9^* mice. This lentiviral pool includes four Inert sgRNAs, as well as sgRNAs targeting *Lkb1*, *Rb1*, and *Rbm10* as tumor suppressor controls. Each candidate gene from the initial screen is targeted with three sgRNAs. Tumors were initiated through intratracheal delivery of Lenti-sgValidation/Cre, and Tuba-seq was performed on each tumor-bearing lung 12 weeks after initiation, followed by analyses of sgID-BC sequencing data to characterize the effects of inactivating each gene (**b**). **c.** Fluorescence images of representative lung lobes 12 weeks after tumor initiation. Scale bars = 5 mm. Lung lobes are outlined with a white dashed line. **d.** Tumor sizes at indicated percentiles for each sgRNA relative to the size of sgInert-containing tumors at the corresponding percentiles in *KT;H11^LSL-Cas9^* mice. Genes are ordered by 95^th^ percentile tumor size, with sgInerts on the left. Note that sg*Lkb1* is plotted on a separate scale to facilitate visualization of sgRNAs with lesser magnitudes of effect. Dashed line indicates no effect relative to sgInert. Error bars indicate 95% confidence intervals. 95% confidence intervals and P-values were calculated by bootstrap resampling. Percentiles that are significantly different from sgInert (2-sided FDR-adjusted p < 0.05) are in color. **e.** Targeting *Hras* and *Nras* significantly increases mean tumor size relative to sgInerts, assuming a log-normal distribution of tumor sizes (LNmean). Error bars indicate 95% confidence intervals calculated by bootstrap resampling. **f.** Schematic of tumor initiation with individual Lenti-sgRNA/Cre vectors. Mouse number and titer of the lentiviral vectors are indicated. **g.** Representative fluorescence images of lungs from *KT;H11^LSL-Cas9^* mice after tumor initiation with Lenti-sgRNA/Cre vectors as indicated. Scale bar = 5 mm. **h.** Representative H&E images of lungs from *KT;H11^LSL-Cas9^* mice after tumor initiation with Lenti-sgRNA/Cre vectors as indicated. Tumor area (percentage of total lung area) from each mouse is shown as Mean ± SD. *: p<0.05; Scale bar = 5 mm. **i.** Tumor burden in *KT;H11^LSL-Cas9^* mice with tumors initiated with Lenti-sgRNA/Cre vectors as indicated. Each dot represents relative tumor area (percentage of total lung area) from one mouse. *: p<0.05 **j.** Representative BrdU staining images of lungs from *KT;H11^LSL-Cas9^* mice after tumor initiation with Lenti-sgRNA/Cre vectors as indicated. Number of Brdu^pos^ cells per field is shown as Mean ± SD. **: p<0.01; Scale bar = 100 μm. **k.** Quantification of proliferation cells in *KT;H11^LSL-Cas9^* mice with tumors initiated with Lenti-sgRNA/Cre vectors as indicated. Each dot represents a tumor. **: p<0.01

In addition, we validated the tumor-suppressive function of HRAS and NRAS in oncogenic KRAS-driven lung tumor growth by initiating tumors in *KT;H11^LSL-Cas9^* mice with individual sg*Inert-*, sg*Hras-* and sg*Nras*-containing Lenti-sgRNA/Cre vectors (**Figure 2f**). Inactivation of either *Hras* or *Nras* increased tumor growth as assessed by direct fluorescence and histological analyses (**Figure 2g-k**). Collectively, these results suggest that RAS paralogs constrain the growth of oncogenic KRAS^G12D^-driven lung cancer growth.

### HRAS and NRAS can be growth-suppressive in human lung cancer cells

To assess the relevance of HRAS and NRAS as tumor suppressors in human lung cancer, we tested the function of HRAS and NRAS in oncogenic KRAS-driven human lung adenocarcinoma cell lines. Previous genome-scale CRISPR/Cas9 screens revealed that inactivating these genes was generally detrimental to cancer cell line growth under standard culture conditions (**Figure S5a)**^10, 35^. Interestingly, HRAS and NRAS suppressed the growth of oncogenic KRAS^G12S^-driven A549 cells and of several oncogenic KRAS-driven lung cancer cell lines when grown in 3D culture conditions, suggesting that these genes can function as tumor suppressors in certain contexts (**Figure S5b-c**)^10, 15^. To further assess the functions of HRAS and NRAS in oncogenic KRAS-driven human adenocarcinoma cell lines, we performed gain and loss of function studies on H23 (KRAS^G12C/+^) and H727 (KRAS^G12V/+^) cells under growth factor restricted growth conditions. We inactivated *HRAS* and *NRAS* using CRISPR/Cas9 and generated variants with doxycycline-inducible wild-type *HRAS* re-expression. Inactivation of *HRAS* or *NRAS* in oncogenic KRAS-driven cells increased cell growth when cells were grown with limited serum and increased clonal growth potential when cells were grown in anchorage-independent conditions (**Figure 3a, c, d**). Conversely, re-expression of HRAS in these *HRAS*-null cells impaired proliferation and clonal growth (**Figure 3b, e, f**). H23 cells with inactivated *HRAS* or *NRAS* also formed larger and more proliferative tumors after intravenous and subcutaneous transplantation (**Figure 3g-k, Figure S6**). These results demonstrate that wild-type HRAS and NRAS can also function as tumor suppressors in oncogenic KRAS-driven human lung cancer cells *in vitro* and *in vivo*. This consistency between human cell culture and autochthonous mouse models further suggests that HRAS and NRAS are tumor suppressors in oncogenic KRAS-driven lung adenocarcinoma.

**Figure 3.**
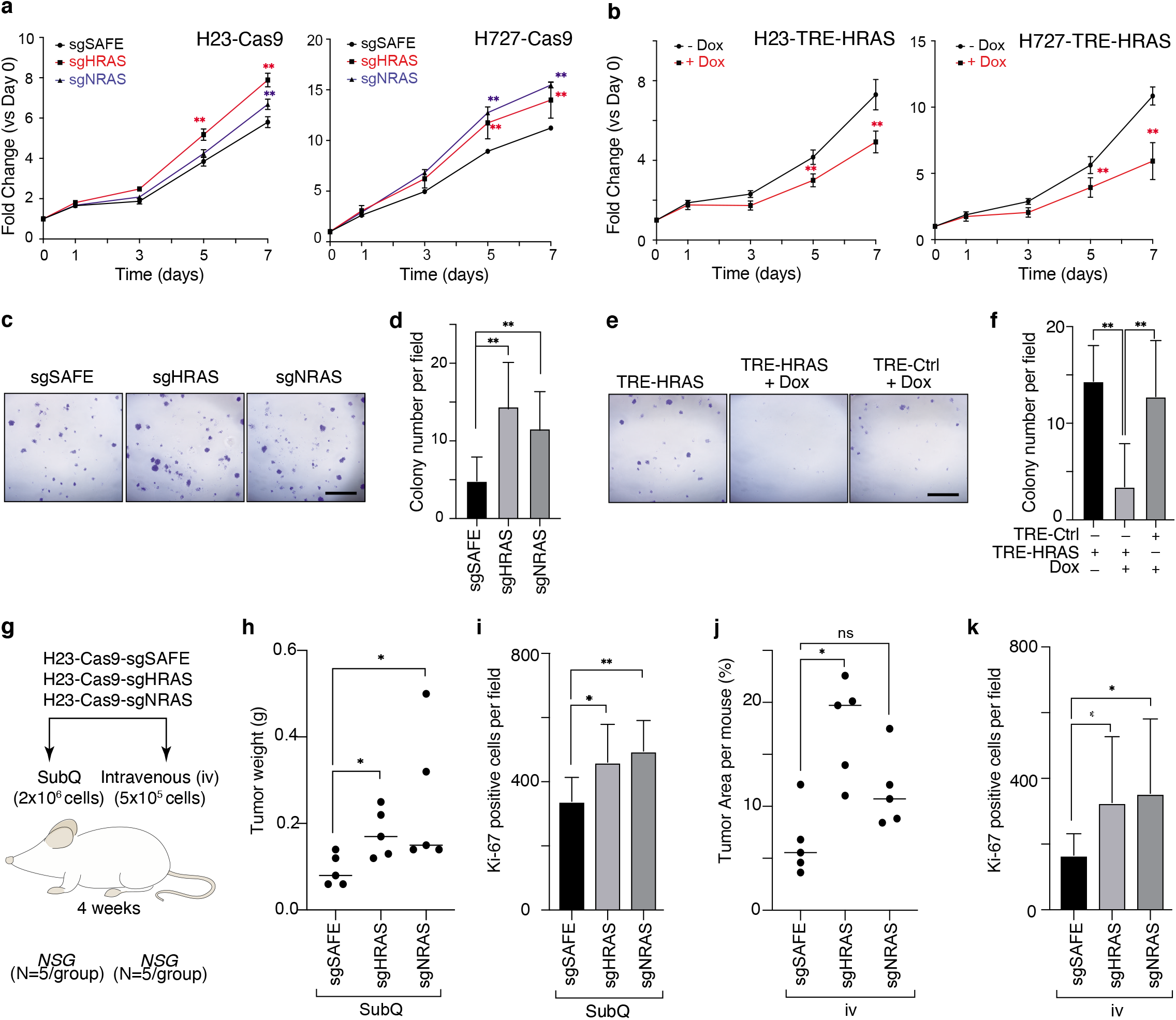
Wildtype HRAS or NRAS constrain the growth of human KRAS-driven cancer cell lines. **a.** Inactivation of wild type HRAS or NRAS increases growth of KRAS-mutant H23 (G12C) and H727 (G12V) cells. Wildtype (sg*SAFE*) or *HRAS*- or *NRAS*-knockout cells were seeded in 96 well plates and cultured under limited serum (1%). Cell numbers were measured via CCK8 assay. Points are Mean±SD of 12 wells normalized to Day 0. **: p<0.01 **b.** Re-expression of wild type HRAS suppresses proliferation of HRAS-null H23 and H727 cells. TRE-HRAS cells were seeded in 96 well plates and cultured under limited serum (1%) with or without 50 ng/ml Doxycycline (Dox) and cell numbers were measured via CCK8 assay. Points are Mean±SD of 12 wells normalized to Day 0. **: p<0.01 **c-d.** Inactivation of HRAS or NRAS increases H23 colony formation. Wildtype (sg*SAFE*), HRAS-knockout (sg*HRAS*), or NRAS-knockout (sg*NRAS*) H23 cells were seeded at 1000 cells/well in 6-well plates and grown for two weeks. Cells were stained with crystal violet. **c.** Representative images. Scale bar = 5mm. d. Mean±SD of colony number of 12 fields. **: p<0.01 **e-f.** Re-expression of wild type HRAS suppresses HRAS-null H23 cell colony formation. TRE-Ctrl or TRE-HRAS H23 cells were seeded at 1000 cells/well in 6-well plates and grown with or without 50 ng/ml Dox for two weeks. Cells were stained with crystal violet. **e.** Representative images. Scale bar = 5mm. f. Mean±SD of colony number of 12 fields. **: p<0.01 **g-k.** Inactivation of wild type HRAS or NRAS increases H23 cell growth after transplantation. **g.** Schematic of tumor initiation with subcutaneous (SubQ) or intravenous (IV) transplantation of H23 cells with inactivation of HRAS or NRAS in NSG mice. Mouse number, cell number, and tumor growth time after transplantation are indicated. **h.** Tumor weight from SubQ transplantation of indicated cells. Each dot represents a mouse. Mean value was shown. **i.** Ki67^pos^ cell number in tumor section from SubQ transplantation of indicated cells was shown as Mean±SD value of 20 view fields. **j.** Tumor area (percentage of h-mitochondriapos area) from IV transplantation of indicated cells. Each dot represents a tumor. Mean value was shown. **k.** Ki67^pos^ cell number in tumor section from IV transplantation of indicated cells is shown as Mean±SD value of 20 view fields (200x). *: p<0.05; **: p<0.01; ns: not significant.

### Inactivation of RAS paralogs increases signaling downstream of oncogenic KRAS

Wild-type KRAS has been shown to be tumor-suppressive in multiple experimental models of oncogenic KRAS-driven cancer, likely due to its ability to interact with and antagonize oncogenic KRAS^36–38^. We have demonstrated that wild-type HRAS and NRAS suppress oncogenic KRAS^G12D^-driven lung cancer growth *in vivo*. Thus, to further explore the molecular mechanism driving this effect, we initially assessed whether HRAS and NRAS alter signaling downstream of oncogenic KRAS. We initially performed pERK immunohistochemistry on lung tumors initiated with Lenti-sgRNA/Cre vectors containing sg*Inert*, sg*Hras* or sg*Nras* in *KT;H11^LSL-Cas^*^9^ mice. Inactivation of HRAS or NRAS increased the number of pERK-positive cells in KRAS^G12D^-driven lung cancer (**Figure 4a, Figure S7a**). Subcutaneous tumors from transplanted H23 cells with inactivated *HRAS* or *NRAS* also contained more pERK-positive cells when compared to tumors from transplanted wildtype (sg*SAFE*) H23 cells (**Figure 4b, Figure S7b**). In addition, sorted cancer cells from *KT;H11^LSL-Cas9^* mice with lung tumors initiated with Lenti-sg*Hras*/Cre also had greater pERK and pAKT compared to those from tumors initiated with Lenti-sg*Inert*/Cre (**Figure 4c, Figure S7c**). Inactivation of either *Hras* or *Nras* in mouse (HC494) or human (H23 and HOP62) oncogenic KRAS-driven cell lines increased ERK phosphorylation, while their effects on AKT phosphorylation was more cell context dependent (**Figure 4d-e, Figure S7d-e**). Conversely, re-expression of wild-type HRAS in HRAS-null H23 and HOP62 human lung cancer cells reduced ERK phosphorylation again with cell context dependent effect on AKT phosphorylation (**Figure 4f, Figure S7f**). Previous publications have shown that inactivating wild-type KRAS increases sensitivity to MEK inhibitors^37, 39^. Consistent with these studies, we found that inactivation of *HRAS* in H23 cells increased sensitivity to the MEK inhibitor trametinib while re-expression of HRAS made cells more resistant (**Figure 4g, h**). These data suggest that inactivation of *HRAS* or *NRAS* hyper-activates MAPK-ERK signaling in KRAS mutant cancer cells^40–42^.

**Figure 4.**
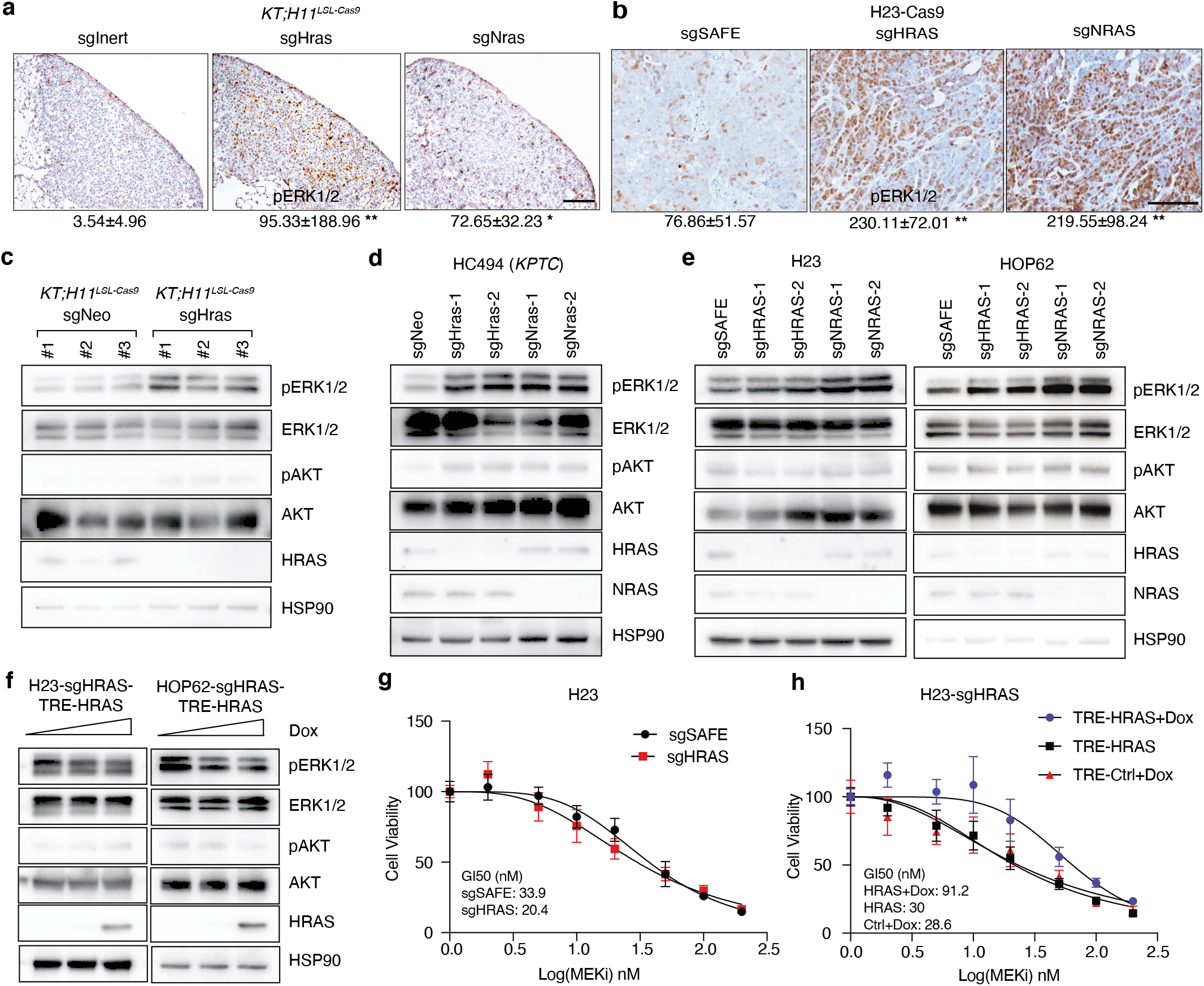
Wildtype RAS paralogs suppress RAS signaling. **a.** Representative image of pERK staining in *KT;H11^LSL-Cas9^* mice with tumors initiated with Lenti-sgRNA/Cre vectors as indicated. Quantification of pERK^pos^ cells per tumor was shown as Mean±SD of 20 tumors. *: p<0.05; **: p<0.01; Scale bar: 100 μm **b.** Representative image of pERK staining in subcutaneous tumor transplanted with H23 cells as indicated. Quantification of pERK^pos^ cells per field was shown as Mean±SD of 20 fields. **: p<0.01; Scale bar: 100 μm. HSP90 shows loading. **c.** Western blot analysis of sorted cancer cells from *KT;H11^LSL-Cas9^* mice transduced with Lenti-sgRNA/Cre vectors as indicated. Multiple tumors were pooled and Tomato^pos^ cancer cells were sorted prior to and protein extraction. HSP90 shows loading. **d.** Western blot analysis of murine lung adenocarcinoma cell line that was transduced with Lenti-sgRNA vectors as indicated and selected with puromycin to generate stable knockout cell lines. Wildtype cells (sg*Neo*) or HRAS- or NRAS-knockout cells (sg*Hras*, sg*Nras*) were cultured under limited serum (1%) for 2 days before protein extraction. HSP90 shows loading. **e.** Western blot analysis of cultured human lung adenocarcinoma cell lines transduced with Lenti-sgRNA vectors as indicated and selected with puromycin to generate stable knockout cell lines. Wildtype cell (sg*SAFE*) or HRAS- or NRAS-knockout cells (sg*HRAS*, sg*NRAS*) were cultured under limited serum (1%) for 2 days before protein extraction. HSP90 shows loading. **f.** Western blot analysis of human lung adenocarcinoma cell lines re-expression HRAS (TRE-HRAS) under Doxycycline (Dox) treatment. HRAS-null cells were generated as described in Figure 3a. HRAS-null cells were re-transduced with lentiviral vector expressing TRE-HRAS at high MOI (>5) to generate stable HRAS re-expression cells (sgHRAS-TRE-HRAS). To re-express HRAS, cells were treated with 0, 1, or 2ng/ml Dox and cultured under limited serum (1%) for 2 days before protein extraction. HSP90 shows loading. **g.** Comparison of GI50 values to MEK inhibitors trametinib among wildtype and HRAS-null H23 cells under treatment of indicated dose of trametinib for four days. Cell numbers were measured via CCK8 assay and normalized to cells treated with vehicle. Each data point was shown as Mean±SD of 12 wells. **h.** Comparison of GI50 values to MEK inhibitors trametinib among HRAS-null H23 cells (H23-sgHRAS) re-expressing HRAS in presence (HRAS+Dox) or absence (HRAS) of Doxycycline plus indicated dose of trametinib for four days. Cell numbers were measured via CCK8 assay and normalized to cells treated with vehicle. Each data point was shown as Mean±SD of 12 wells.

### RAS paralogs suppress oncogenic KRAS-KRAS interaction

RAS proteins interact and form functional clusters on membranes to efficiently recruit downstream effectors^43–45^. Whether RAS proteins form dimers or oligomers through direct interactions or through close physical proximity is debated within the field ^16, 46–48^. We next assessed whether HRAS and NRAS “interact” with KRAS without attempting to distinguish direct from proximity-driven interactions. AP/MS data suggest that all three RAS proteins are able to interact with their paralogs, supporting the existence of heterotypic RAS-RAS interactions (**Figure 5a**). To assess the ability of RAS paralogs to interact with oncogenic KRAS^G12D^, we adapted a luminescent reporter system (ReBiL2.0 system), which relies on luciferase complementation to quantify RAS-RAS interactions in living cells^16^ (**Figure 5b**). Through expression of wild-type KRAS, HRAS, or NRAS in KRAS^G12D^-KRAS^G12D^ interaction reporter cells and control reporter cells, we found that all wild-type RAS paralogs are able to disrupt KRAS^G12D^-KRAS^G12D^ interactions **(Figure 5c, Figure S10a)**.

**Figure 5.**
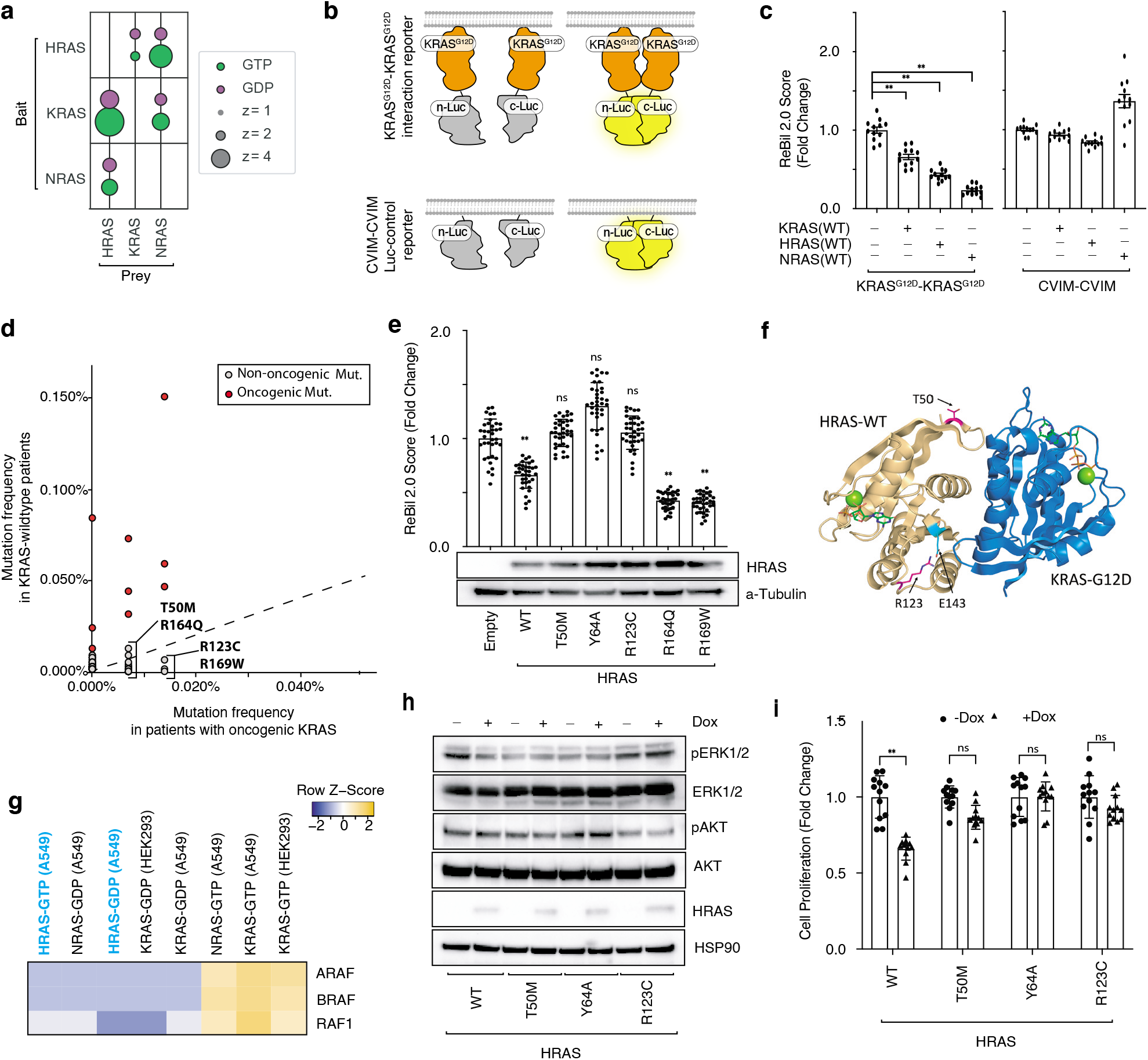
Wildtype RAS paralogs fine-tune RAS signaling through interaction with oncogenic KRAS. **a.** Bubble plot of three AP/MS experiments with H-, K-, and N-RAS as baits (rows), showing the enrichment of their paralogs (columns). **b.** Diagram of the ReBiL2.0 system. KRAS^G12D^-KRAS^G12D^ interactions were quantified by normalized luminescent signal generated by membrane association facilitated interaction of the split-luciferase that is fused to the N-terminus of KRAS^G12D^ (upper). Split-luciferase that is fused to the last four amino acids of KRAS (CVIM) is applied as control for background split-luciferase interaction on the membrane (lower). Adapted from Li et al. 2020. **c.** All three RAS proteins are able to disrupt KRAS^G12D^-KRAS^G12D^ interaction. U2OS-764 (nl-KRAS^G12D^/cl-KRAS^G12D^) or U2OS-794 (nl-CVIM/cl-CVIM) cells expressing KRAS, HRAS, or NRAS were cultured in limited serum (1%) under 100 ng/ml Doxycycline (Dox) for 24 hours. ReBiL2.0 assay were performed as previously described and detailed in Methods. Points are Mean±SD ReBiL2.0 score of 36 wells normalized to cells transduced with empty lentiviral vector. **: p<0.01 **d.** Pan-cancer frequency of HRAS mutations in patients with wildtype and oncogenic KRAS-tumors from Project GENIE. Known oncogenic HRAS mutations are highlighted in red. The dashed line indicates equal mutation frequency in KRAS-wildtype and mutant samples. Four candidate mutations that were chosen for further validation in this study were highlighted. **e.** HRAS^T50M^ and HRAS^R123C^ are novel RAS-RAS interaction deficient mutations. U2OS-764 (nl-KRAS^G12D^/cl-KRAS^G12D^) cells expressing wildtype or rare mutant HRAS were cultured in limited serum (1%) under 100 ng/ml Dox for 24 hours. Points are Mean±SD ReBiL2.0 score of 12 wells normalized to cells transduced with empty lentiviral vector (upper). **: p<0.01; ns: not significant. HRAS (wildtype and mutant) protein expression level in corresponding cells were shown by Western blot analysis (lower). **f.** HRAS^T50M^ and HRAS^R123C^ are located close to the predicted HRAS-KRAS interaction interface. HRAS is shown in light orange and KRAS^G12D^ is shown in blue. Residue R123 (in magenta) makes an intrachain salt bridge with E143 (in cyan). **g.** Prey RAF proteins enriched in each experiment with the indicated baits in A549 cells (for K-, H-, or N-RAS) or HEK293 cells (for KRAS). Yellow color indicates higher values of NSAF. Both GTP- and GDP-bond HRAS behave like GDP-bond KRAS in their RAF interactions. **h.** Western blot analysis of cultured HRAS-null HOP62 cells (HOP62-Cas9-sgHRAS) re-expressing wildtype or mutants (T50M, Y64A, or R123C) under Dox treatment. Cells were cultured under limited serum (1%) for 2 days before protein extraction. Re-expression of HRAS mutations have no effects on ERK phosphorylation. **i.** Cell proliferation of cultured HRAS-null HOP62 cells (HOP62-Cas9-sgHRAS) re-expressing wildtype or mutants (T50M, Y64A, or R123C) under Dox treatment. Cells were cultured in limited serum (1%) with or without Dox for 4 days. Cell viability was measured via CCK8 assay and normalized to cells treated with vehicle. Re-expression of HRAS mutants have no effects on cell proliferation.

### Patient-derived HRAS^T50M^ and HRAS^R123C^ mutations impair interaction of HRAS with oncogenic KRAS and abrogate its tumor suppressive function

Our findings suggest that the tumor-suppressive function of wild-type HRAS is mediated by competitive interactions with oncogenic KRAS, therefore we hypothesized that there could be *HRAS* mutations in human tumors with oncogenic *KRAS* that impair this interaction. To evaluate this possibility, we analyzed data from AACR’s Genomics Evidence Neoplasia Information Exchange (GENIE). Mutations in *HRAS* were rare (pan-cancer frequency of non-synonymous mutations was 1.32%) and about half (0.57%) were oncogenic mutations in codons 12, 13 or 61 that occurred in samples lacking oncogenic *KRAS* (**Figure S8a**). We did, however, identify multiple rare non-oncogenic *HRAS* mutations in oncogenic KRAS containing lung adenocarcinomas and tumors of other types (**Figure 5d, Figure S8**). To test whether these mutants lack the ability to interact with oncogenic KRAS, we used the ReBiL2.0 system. We measured the ability of four of these *HRAS* mutants, as well as a control Y64A mutant that has been suggested to reduce HRAS-HRAS dimerization^47^, to inhibit KRAS^G12D^-KRAS^G12D^ interactions. We identified two HRAS mutants, T50M and R123C, that are unable to reduce KRAS^G12D^-KRAS^G12D^ interactions (**Figure 5e, Figure S10b**). Interestingly, both HRAS^T50^ and HRAS^R123^ are located close to the predicted HRAS-KRAS^G12D^ interface involving the α4 and α5 helices (**Figure 5f, Figure S9**). R123 is involved in an intrachain salt bridge with residue E143, which also participates in the RAS-RAS interface. Mutation to cysteine results in an uncompensated charge on E143, which may destabilize the RAS-RAS interaction. These findings are consistent with a model in which wild-type RAS paralogs competitively interacts with oncogenic KRAS and thus suppress KRAS^G12D^-KRAS^G12D^ interactions.

Previous publications have shown that different RAS proteins preferential bind to RAF proteins and other RAS effectors and thus could function differently in their downstream signaling^10, 50^. Re-analysis of HRAS and NRAS AP/MS datasets suggests that GTP-bound HRAS is more similar in its low binding affinity to RAF effectors as GDP-bound rather than GTP-bound KRAS and NRAS (**Figure 5g**)^10^. To test our hypothesis that the disruption of KRAS^G12D^-KRAS^G12D^ interaction by HRAS suppresses downstream oncogenic signaling, we re-expressed wild-type HRAS, HRAS^Y64A^, or the two novel patient-derived HRAS^T50M^ and HRAS^R123C^ mutants in *HRAS*-null lung cancer cells. Re-expression of wild-type HRAS, but not any of the three mutants, reduced ERK phosphorylation and proliferation (**Figure 5h-i, Figure S10c**). These results further suggest that RAS paralog imbalance alters oncogenic KRAS signaling via oncogenic KRAS-wildtype RAS paralog interaction and thus is a driver of lung cancer growth.

### HRAS and NRAS are specific suppressors of oncogenic KRAS-driven lung cancer growth

Our *in vivo* data demonstrate that HRAS and NRAS function as tumor suppressors, and our cell culture results suggest that these effects may be mediated by interaction of these RAS paralogs with oncogenic KRAS. If the tumor-suppressive mechanism by which HRAS and NRAS is mediated through interactions with oncogenic KRAS, then these genes should not be tumor suppressors in lung adenocarcinoma in which activation of the RAS/RAF/MEK signaling pathway occurs downstream of KRAS. To test this directly in autochthonous tumors, we initiated tumors with a sub-pool of barcoded lenti-sgRNA/Cre vectors (Lenti-sgMultiGEMM/Cre) in mouse models of oncogenic KRAS-driven and oncogenic BRAF-driven lung cancer (**Figure 6a**). In addition to vectors targeting *Hras* and *Nras*, this pool contained vectors targeting several known tumor suppressors (*Apc, Rbm10*, and *Cdkn2a*) and other KRAS-interacting proteins (*Aldh1a, Nme2*), as well as control vectors (**Figure 6a**). We initiated tumors with the Lenti-sgMultiGEMM/Cre pool in *KT* and *KT;H11^LSL-Cas^*^9^ mice as well as in *BrafT;H11^LSL-Cas9^* mice which contain a Cre-regulated allele of oncogenic BRAF^V618E^ (the mouse equivalent of BRAF^V600E^)(**Figure 6b**)^51^. 15 weeks after tumor initiation *BrafT;H11^LSL-Cas9^* mice has greater overall tumor burden than *KT;H11^LSL-Cas9^* mice (**Figure 6c-d**). Analysis of the distribution of sgInert tumor sizes in the two models using Tuba-seq showed that oncogenic BRAF-driven tumors were larger than oncogenic KRAS-driven tumors (median sizes of ∼3500 cells and ∼1000 cells, respectively). The two distributions had similar maximum tumor sizes, suggesting that the increased tumor burden is driven by a shift towards larger tumors of relatively uniform size which is consistent with previous results (**Figure 6e-f**)^51^.

**Figure 6.**
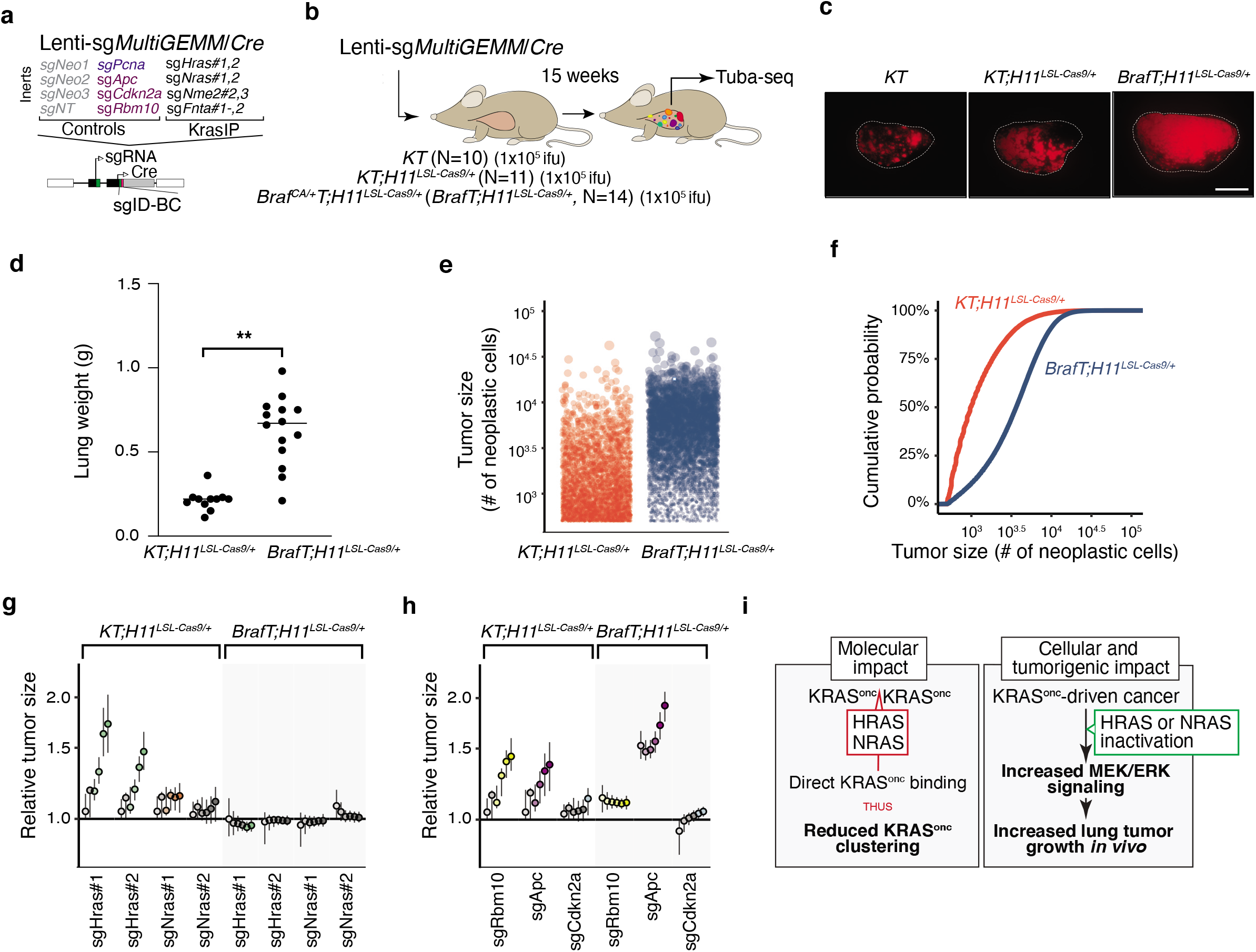
Paired screens in KRAS-driven and BRAF-driven lung cancer models validates HRAS and NRAS as KRAS-specific tumor suppressors. **a-b.** Schematic of pairwise screen of tumor suppressive function in KRAS- and BRAF-driven lung cancer. A pool of barcoded Lenti-sgRNA/Cre vectors targeting top mediators of KRAS-driven lung tumor growth (Lenti-sgMultiGEMM/Cre) was used to initiate tumors in cohorts of *KT;H11^LSL-Cas9/+^* and *Braf^CA/+^T;H11^LSL-Cas9 /+^* (*BrafT;H11^LSL-Cas9/+^*) mice. Each regulator of KRAS-driven tumor growth (*Hras*, *Nras*, *Nme2* and *Fnta*) was targeted by two sgRNAs (those with the largest effect size in the validation screen). The pool also included four Inert sgRNAs, as well as sgRNAs targeting *Apc*, *Cdkn2a*, and *Rbm10* as tumor suppressor controls (**a**). Tumors were initiated through intratracheal delivery of Lenti-sgMultiGEMM/Cre, and Tuba-seq was performed on each tumor-bearing lung 15 weeks after initiation, followed by analysis of sgID-BC sequencing data to characterize the effects of inactivating each gene (**b**). **c.** Fluorescence images of representative lung lobes 15 weeks after tumor initiation. Scale bars = 5 mm. Lung lobes are outlined with a white dashed line. **d.** Total lung weight in *KT;H11^LSL-Cas9/+^* and *BrafT;H11^LSL-Cas9/+^* mice 15 weeks after tumor initiation. Each dot is a mouse and mean value is indicated. **: p<0.01 **e-f.** Size distribution of sgInert tumors in *KT;H11^LSL-Cas9/+^* and *BrafT;H11^LSL-Cas9/+^* mice. In **e**., each dot represents a tumor, and the area of each dot is proportional to the number of cancer cells in that tumor. To prevent overplotting a random sample of 1,000 tumors from each of five representative *KT;H11^LSL-Cas9/+^* and *BrafT;H11^LSL-Cas9/+^* mice are plotted. In **f.**, the empirical cumulative distribution function of tumor sizes across all *KT;H11^LSL-Cas9/+^* and *BrafT;H11^LSL-Cas9/+^* mice are plotted. Tumors >500 cells in size are shown. **g.** Inactivation of *Hras* and *Nras* increases tumor size in *KT;H11^LSL-Cas9/+^* but not *BrafT;H11^LSL-Cas9/+^* models. Tumor sizes at indicated percentiles for each sgRNA relative to the size of sgInert-containing tumors at the corresponding percentiles in *KT;H11^LSL-Cas9/+^* (left, white background) and *BrafT;H11^LSL-Cas9/+^* (right, gray background) mice. Line at y=1 indicates no effect relative to sgInert. Error bars indicate 95% confidence intervals. Percentiles that are significantly different from sgInert (two-sided FDR-adjusted p < 0.05) are in color. Confidence intervals and P-values were calculated by bootstrap resampling. **h.** Comparison of the effects of inactivation of known tumor suppressors (*Rbm10*, *Apc*, and *Cdkn2a*) on tumor size in *KT;H11^LSL-Cas9/+^* and *BrafT;H11^LSL-Cas9/+^* models. Tumor sizes at indicated percentiles for each sgRNA relative to the size of sgInert-containing tumors at the corresponding percentiles in *KT;H11^LSL-Cas9/+^* (left, white background) and *BrafT;H11^LSL-Cas9/+^* (right, gray background) mice. Line at y=1 indicates no effect relative to sgInert. Error bars indicate 95% confidence intervals. Percentiles that are significantly different from sgInert (two-sided FDR-adjusted p < 0.05) are in color. Confidence intervals and P-values were calculated by bootstrap resampling. **i.** Wildtype RAS paralogs function as tumor suppressors in oncogenic KRAS-driven lung cancer. Left panel, in oncogenic KRAS-driven lung cancer cells, wildtype RAS paralogs competitively interact with oncogenic KRAS and suppress oncogenic KRAS clustering. Right panel, inactivation of wildtype RAS allele, or “RAS paralog imbalance”, hyper-activate oncogenic KRAS signaling and promotes lung cancer growth.

Our Tuba-seq data also allowed us to compare the impact of the CRISRP/Cas9 inactivated genes across oncogenic contexts. Importantly, while inactivation of *Hras* or *Nras* increased the growth of oncogenic KRAS-driven lung tumors, inactivation of *Hras* or *Nras* had no effect on the growth of oncogenic BRAF-driven lung cancer (**Figure 6g, Figure S11d-e**). These results were consistent for both Lenti-sgRNA/Cre vectors targeted each gene. The known tumor suppressor genes assayed (*Apc*, *Cdkn2a*, and *Rbm10*) generally retained their growth-suppressive effects in the BRAF-driven model, suggesting that the abrogation of effect observed for *Hras* and *Nras* is not due to some generic inability of additional alterations to increase BRAF-driven lung tumor growth (**Figure 6h, Figure S11d-e**). Thus, HRAS and NRAS function as specific suppressors of oncogenic KRAS-driven tumor growth *in vivo*.

Assessing the impact of genomic alterations on the growth of lung cancer driven by distinct oncogenes was illuminating in two other regards. First, we identify instances of oncogene-tumor suppressor epistasis (e.g., *Apc* inactivation has a greater effect on BRAF-driven lung cancer whereas *Rbm10* inactivation has a greater effect on KRAS-driven lung cancer) (**Figure 6h, Figure S11d-e**). Thus, the consequences of inactivating tumor suppressor pathways can depend on the oncogenic context. Second, inactivation of *Nme2*, *Fnta*, and *Aldh1a* reduced initiation and growth of both oncogenic KRAS-driven and oncogenic BRAF-driven lung cancer, suggesting that they are generally required for optimal lung cancer growth *in vivo* (**Figure S11**). Thus, our paired screens not only localized the effect of *Hras* and *Nras* inactivation, but also highlighted the value of this approach in uncovering alterations that have effects within or across oncogenic contexts.

## DISCUSSION

Oncogenic KRAS-driven lung cancer is a leading cause of cancer-related deaths. However, despite the identification of oncogenic RAS almost half a century ago, the functions of many RAS-interacting proteins remain largely unknown. By integrating AP/MS data from human cancer cells with somatic cell CRISPR/Cas9-editing in autochthonous mouse models, we assess the functional impact of inactivating a panel of KRAS-interacting proteins on lung cancer *in vivo* in a multiplexed manner. Our results support a model in which heterotypic interactions between RAS paralogs suppress oncogenic KRAS-driven lung cancer growth.

All RAS family proteins, HRAS, NRAS and KRAS (including both the KRAS4A and KRAS4B splice isoforms), have been reported to form dimers and nanoclusters^16, 46–48^. Importantly, both *in vitro* and *in vivo* studies suggest that KRAS-KRAS interactions are required for effector protein activation, cellular transformation, and optimal tumor growth^45^. Furthermore, oncogenic KRAS-wild-type KRAS interactions influence lung cancer initiation, progression, and therapeutic sensitivity^37^. Multiple lines of evidence, including oncogenic *KRAS* copy number gain and loss of the wild-type *KRAS* allele in human tumors, as well as functional studies in mouse models, suggest that wild-type KRAS is tumor-suppressive (also called “RAS allelic imbalance”), although the exact role of wild-type KRAS in lung cancer is still debated^3, 38, 41, 52, 53^. Recent data also suggest that interactions among H-, N- and KRAS occur, thus raising the question of the roles of wild-type HRAS and NRAS in oncogenic KRAS-driven cancer^10, 11, 16^.

In this study, we identified wild-type HRAS and NRAS as potent KRAS-specific tumor suppressors that interact with oncogenic KRAS, disrupt KRAS-KRAS interactions, and suppress RAS/MAPK signaling. Inactivation of *HRAS* or *NRAS* in the context of oncogenic KRAS led to an increase in downstream MAPK signaling (**Figure 4**). The impact of RAS paralog imbalance extends beyond lung cancer and KRAS codon 12 mutations. Germline Hras deletion increases the development of Kras-driven pancreatic cancer, skin papilloma, and carcinogen induced KRAS^Q61^ lung cancer^53–55^. Interestingly, we also identified two rare, patient-derived *HRAS* mutations, HRAS^T50M^ and HRAS^R123C^, which are incapable of disrupting KRAS clustering, and would therefore likely confer fitness advantages to oncogenic KRAS-driven cancer. These results suggest that modulating RAS protein interactions, such as by skewing the stoichiometry of oncogenic to wild-type RAS or forcing inter-paralog competition, could lead to novel therapeutic strategies. However, the dynamics of intracellular RAS interactions, as well as the importance of these mutations in oncogenesis requires further study.

Given the complexity of RAS signaling, other non-mutually exclusive mechanisms by which RAS paralogs could reduce oncogenic KRAS-driven cancer growth should be considered. For example, it has been reported that upstream regulators, such as SOS1, could bridge the interaction between oncogenic and wild-type RAS^56^. GDP-bound wild-type HRAS and NRAS could also compete with oncogenic KRAS for upstream guanine nucleotide exchange factors and thus reduce RAS signaling^57^. In addition, although we provide evidence that inactivation of *Hras* and *Nras* has no impact on oncogenic BRAF-driven lung cancer, it is possible that they could compete with oncogenic KRAS for other BRAF-independent downstream effectors. Whether HRAS and NRAS also function through these alternative routes, and how different mechanisms are synchronized to execute their tumor-suppressive functions, will require additional investigation.

The National Cancer Institute “RAS Pathway V2.0”, contains more than 200 proteins known or suspected to be involved in RAS signaling. Characterizing the role of these proteins in tractable *in vivo* models of RAS-driven cancer remains a challenge. Our study outlines a technological avenue to study KRAS-specific signaling components in a multiplexed manner. By harnessing the power of Tuba-seq, we were able to quantify the tumor suppressive and promoting effects of more than a dozen putative RAS pathway genes simultaneously, highlighting the function of HRAS and NRAS as tumor suppressors. Furthermore, by performing paired screens in oncogenic KRAS-driven and oncogenic BRAF-driven mouse lung cancer models, we localized the growth suppressive effects of these RAS paralogs to lung cancer driven specifically by oncogenic KRAS. Our study thus demonstrates the feasibility of performing *in vivo* genetic interaction screening, and the power of such an approach to provide insight into the mechanisms of tumor suppression. Future studies of this type should enable a more quantitative understanding of the role of RAS pathway components in RAS-driven oncogenicity.

---

**Supplemental Figure 1.**
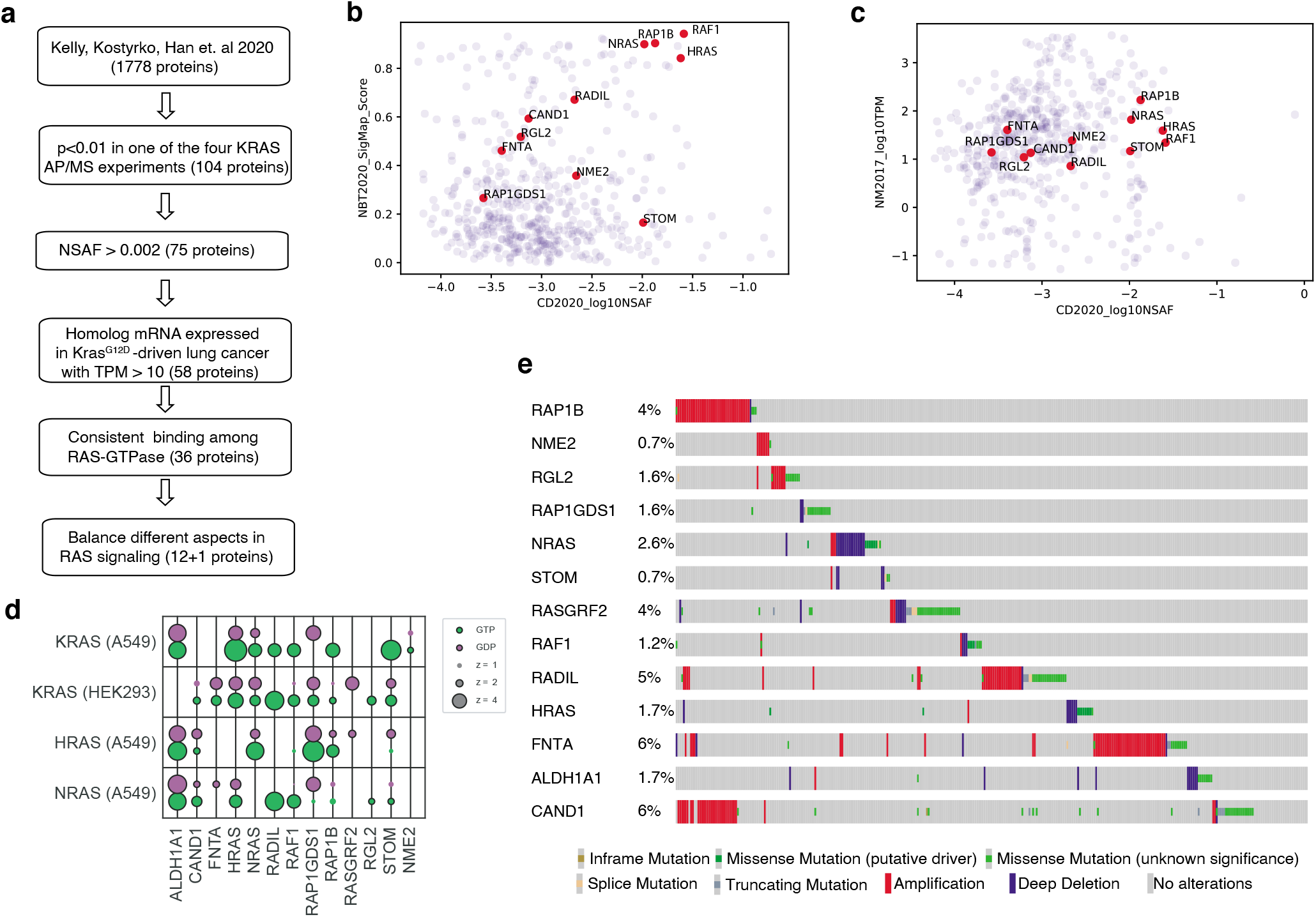
Prioritize candidate KRAS-interacting proteins for this study. **a.** Flow chart for prioritizing candidate KRAS-interacting proteins for this study. Candidate KRAS-interacting proteins were chosen based on multiple criteria including their interaction with KRAS, their homolog mRNA expression in Kras^G12D^-driven lung cancer in mouse model, and the consistency for them to bind different RAS-GTPase. RADIL is added at the last step due to its validated importance in KRAS-mutant human cell lines. **b.** Candidate proteins interact with KRAS from two protein-protein interaction analyses (Kelly, Kostyrko, Han et al. 2020; Broyde, Simpson, Murray et al. 2020). Shared KRAS-interaction proteins are shown as their log10NSAF and SigMap Score. **c.** Homolog mRNA expression (TPM) of candidate KRAS-interacting proteins in Kras^G12D^-driven lung cancer in mouse model (Chuang et al. 2017). **d.** Bubble plot of eight AP/MS experiments with GTP- and GDP-locked mutant GTPases as baits (rows), showing the enrichment of selected candidate KRAS-interacting proteins (columns). Dark borders indicate FDR < 0.05. **e.** Mutation frequencies of these 13 candidate genes in lung adenocarcinoma (data from TCGA, Nat. Genet. 2016).

**Supplemental Figure 2.**
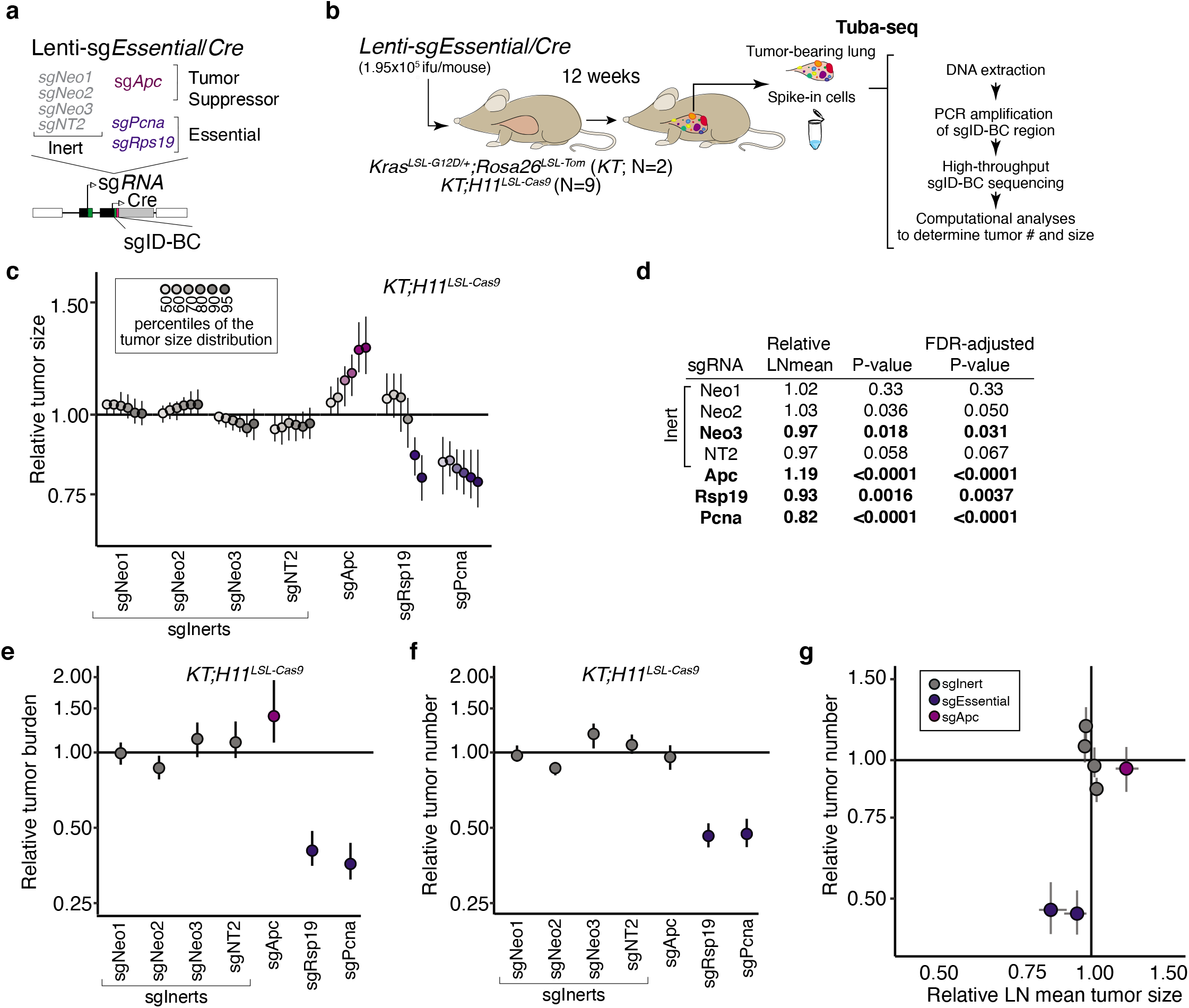
Tumor barcoding coupled with barcode sequencing (Tuba-seq) can uncover engineered alterations that reduce tumor number and growth. **a-b.** Schematic of the Tuba-seq approach to measure the effects of essential gene inactivation on tumor growth. Lentiviral-sgRNA/Cre vectors with inert sgRNAs (gray) or sgRNAs targeting known essential genes (navy) were diversified with a two component sgID-BC. A vector targeting known tumor suppressor *Apc* was included as a positive control (**a**). Tumors were initiated with this barcoded Lenti-sgEssential/Cre pool in *KT* and *KT;H11^LSL-Cas9^* mice. Tuba-seq was performed on each tumor-bearing lung 12 weeks after initiation, followed by analyses of sgID-BC sequencing data to characterize the effects of inactivating each gene (**b**). **c.** Tumor sizes at indicated percentiles for each sgRNA relative to the size of sgInert-containing tumors at the corresponding percentiles. Line at y=1 indicates no effect relative to sgInert. Error bars indicate 95% confidence intervals. Percentiles that are significantly different from sgInert (two-sided FDR-corrected p < 0.05) are in color. Confidence intervals and P-values were calculated by bootstrap resampling. **d.** The impact of each sgRNA on mean tumor size relative to sgInerts, assuming a log-normal distribution of tumor sizes (LNmean). sgRNAs with two-sided P<0.05 after FDR-adjustment are in bold. **e.** The impact of each sgRNA on tumor burden (number of neoplastic cells aggregated across all tumors of a genotype) relative to sgInerts and normalized to the same statistic in *KT* mice to account for representation of each sgRNA in the viral pool. sgInerts are in gray and the line at y=1 indicates no effect. Error bars indicate 95% confidence intervals. Relative burdens significantly different from sgInert (two-sided FDR-corrected p<0.05) are in color. Confidence intervals and P-values were calculated by bootstrap resampling. **f.** The impact of each sgRNA on tumor number relative to sgInerts and normalized to the same statistic in *KT* mice to account for representation of each sgRNA in the viral pool. sgInerts are in gray and the line at y=1 indicates no effect. Error bars indicate 95% confidence intervals. Relative tumor numbers significantly different from sgInert (two-sided FDR-corrected p<0.05) are in color. Confidence intervals and P-values were calculated by bootstrap resampling. **g.** The impact of each sgRNA on tumor number plotted against its impact on LNmean tumor size. The lines at y=1 and x=1 indicate no effect relative to sgInert on tumor number and size, respectively. sg*Rsp19* and sg*Pcna* cluster in the lower left quadrant near x=1, indicating that targeting essential genes strongly reduces tumor number but only moderately decreases average tumor size. Error bars indicate 95% confidence intervals calculated by bootstrap resampling.

**Supplemental Figure 3.**
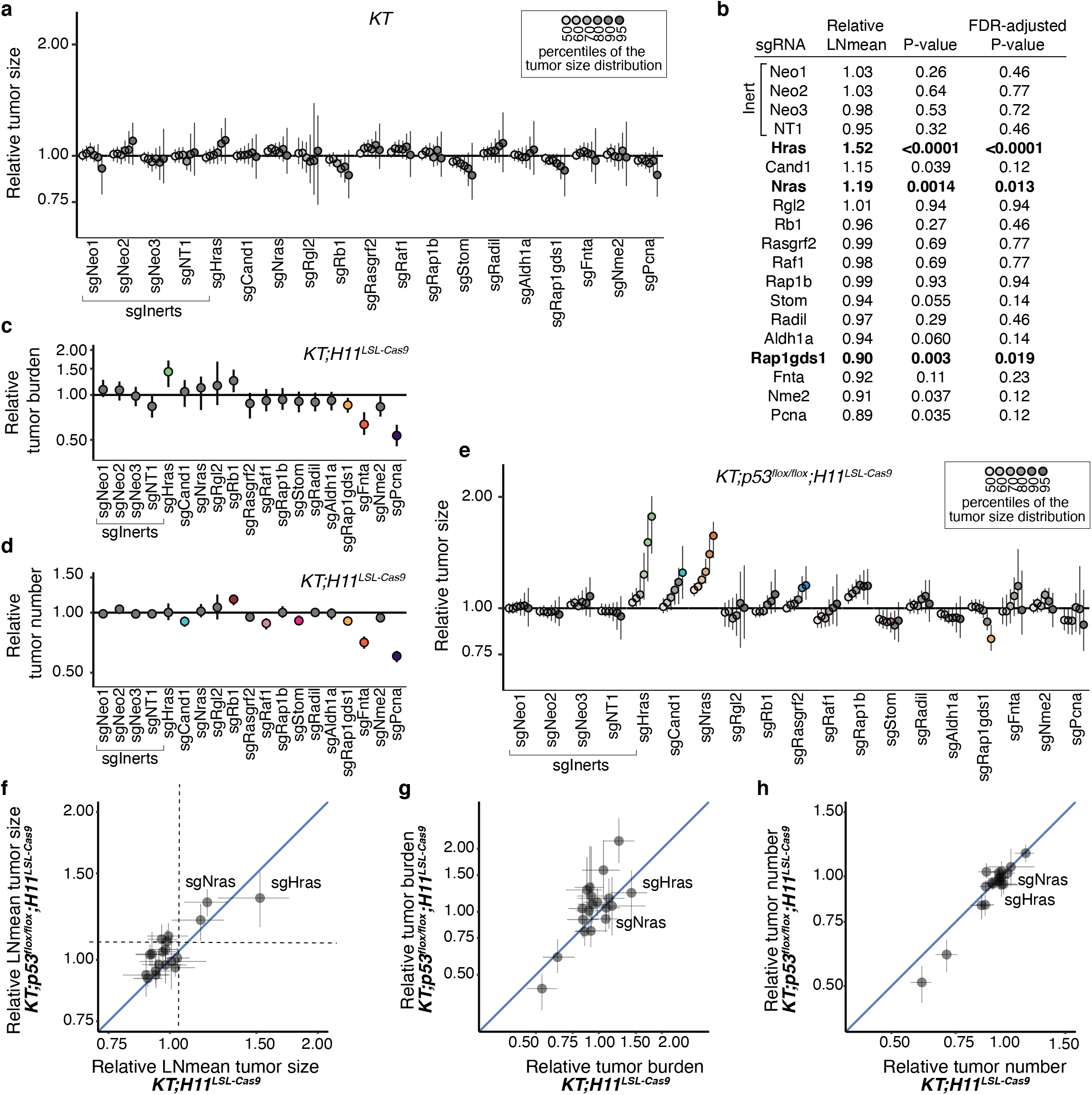
Inactivation of KRAS-interacting proteins has similar impacts on tumor growth in p53-proficient and p53-deficient contexts. **a.** Tumor sizes at indicated percentiles for each sgRNA relative to the size of sgInert-containing tumors at the corresponding percentiles in *KT* mice. *KT* mice lack Cas9, thus all sgRNAs are functionally equivalent to sgInerts. Genes are ordered as in **Figure 1f**. Line at y=1 indicates no effect relative to sgInert. Error bars indicate 95% confidence intervals. Confidence intervals and P-values were calculated by bootstrap resampling. As expected, no percentiles were significantly different from sgInert (two-sided FDR-adjusted p < 0.05). **b.** The impact of each sgRNA on mean tumor size relative to sgInerts in *KT;H11^LSL-Cas9^*, assuming a log-normal distribution of tumor sizes (LNmean). sgRNAs with two-sided P<0.05 after FDR-adjustment are in bold. P-values were calculated by bootstrap resampling. **c-d.** The impact of each sgRNA on tumor burden (**c**) and number (**d**) relative to sgInerts in *KT;H11^LSL-Cas9^* mice, normalized to the corresponding statistic in *KT* mice to account for representation of each sgRNA in the viral pool. sgInerts are in gray and the line at y=1 indicates no effect. Error bars indicate 95% confidence intervals. Relative tumor burdens and numbers significantly different from sgInert (two-sided FDR-adjusted p<0.05) are in color. Confidence intervals and P-values were calculated by bootstrap resampling. **e.** Tumor sizes at the indicated percentiles for each sgRNA relative to the size of sgInert-containing tumors in *KT;p53^flox/flox^;H11^LSL-Cas9^* mice. Genes are ordered as in **Figure 1f**. Dashed line indicates no effect relative to sgInert. Error bars indicate 95% confidence intervals. Percentiles that are significantly different from sgInert (two-sided FDR-adjusted p < 0.05) are in color. Confidence intervals and P-values calculated by bootstrap resampling. **f-h.** Comparison of the impact of each sgRNA on relative LNmean tumor size (**f**), tumor burden (**g**) and tumor number (**h**) in *KT;H11^LSL-Cas9^* and *KT;p53^flox/flox^;H11^LSL-Cas9^* mice. Error bars indicate 95% confidence intervals calculated by bootstrap resampling.

**Supplemental Figure 4.**
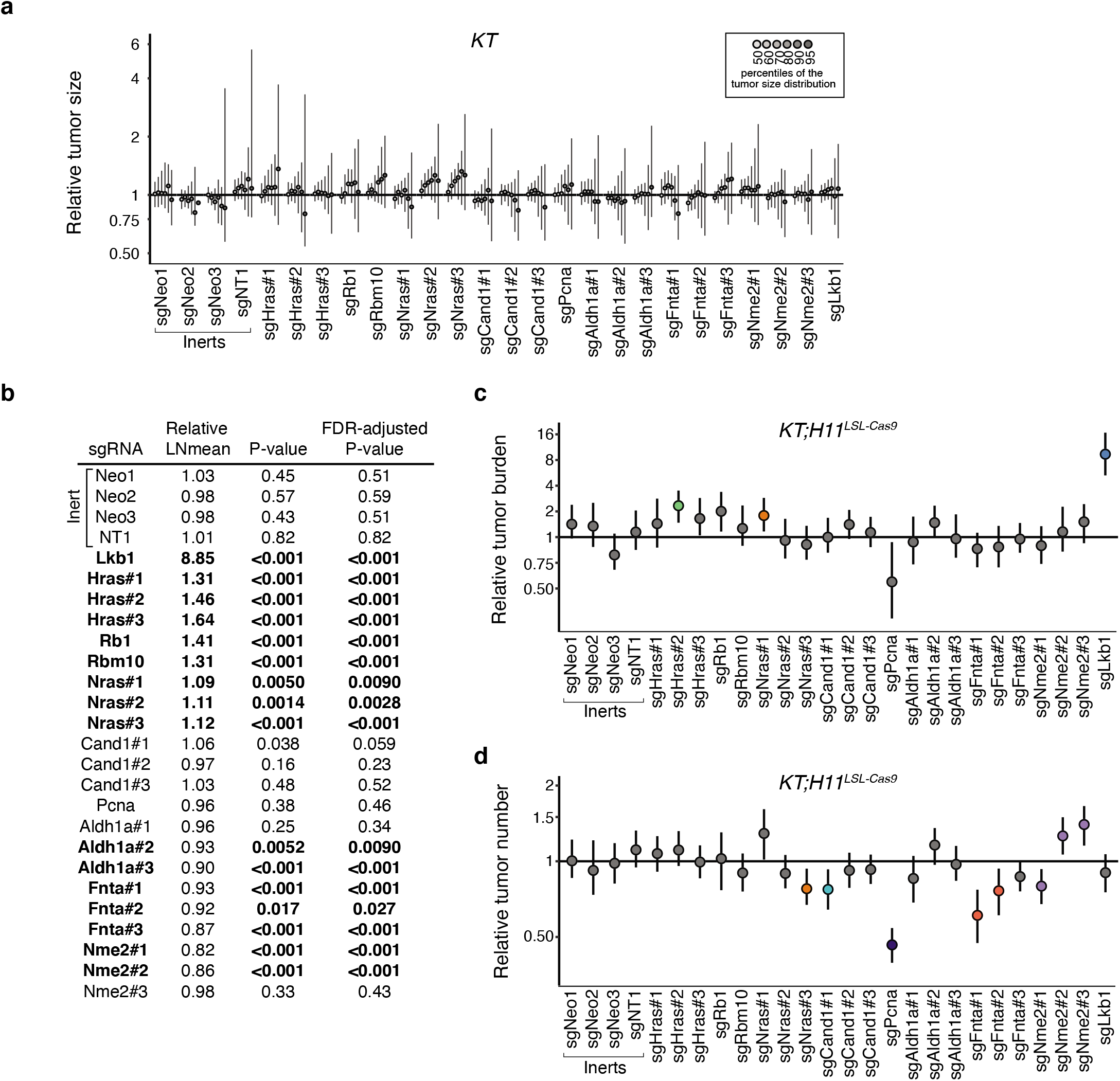
Top candidate KRAS-interacting proteins from initial Tuba-seq screen impact multiple metrics of tumor growth in validation cohort. **a.** Tumor sizes at indicated percentiles for each sgRNA relative to the size of sgInert-containing tumors at the corresponding percentiles in *KT* mice. *KT* mice lack Cas9, thus all sgRNAs are functionally equivalent to sgInerts. Genes are ordered as in **Figure 2d**, but note the change in axis scaling. Line at y=1 indicates no effect relative to sgInerts. Error bars indicate 95% confidence intervals. Confidence intervals and P-values were calculated by bootstrap resampling. As expected, no percentiles were significantly different from sgInert (FDR-adjusted p < 0.05). **b.** The impact of each sgRNA on mean tumor size relative to sgInerts, assuming a log-normal distribution of tumor sizes (LNmean). Two-sided P-values were calculated by bootstrap resampling. sgRNAs with P<0.05 after FDR-adjustment are in bold. Note that this data for the sgInerts, sg*Hra*s#1-3 and sg*Nra*s#1-3 is also plotted in **Figure 2e**. **c.** The impact of each sgRNA on tumor burden relative to sgInerts in *KT;H11^LSL-Cas9^* mice, normalized to the corresponding statistic in *KT* mice to account for representation of each sgRNA in the viral pool. sgInerts are in gray and the line at y=1 indicates no effect. Error bars indicate 95% confidence intervals. Relative tumor burdens significantly different from sgInert (two-sided FDR-adjusted p<0.05) are in color. Confidence intervals and P-values were calculated by bootstrap resampling. **d.** The impact of each sgRNA on tumor number relative to sgInerts in *KT;H11^LSL-Cas9^* mice, normalized to the corresponding statistic in *KT* mice to account for representation of each sgRNA in the viral pool. sgInerts are in gray and the line at y=1 indicates no effect. Error bars indicate 95% confidence intervals. Relative tumor numbers significantly different from sgInert (two-sided FDR-adjusted p<0.05) are in color. Confidence intervals and P-values were calculated by bootstrap resampling.

**Supplemental Figure 5.**
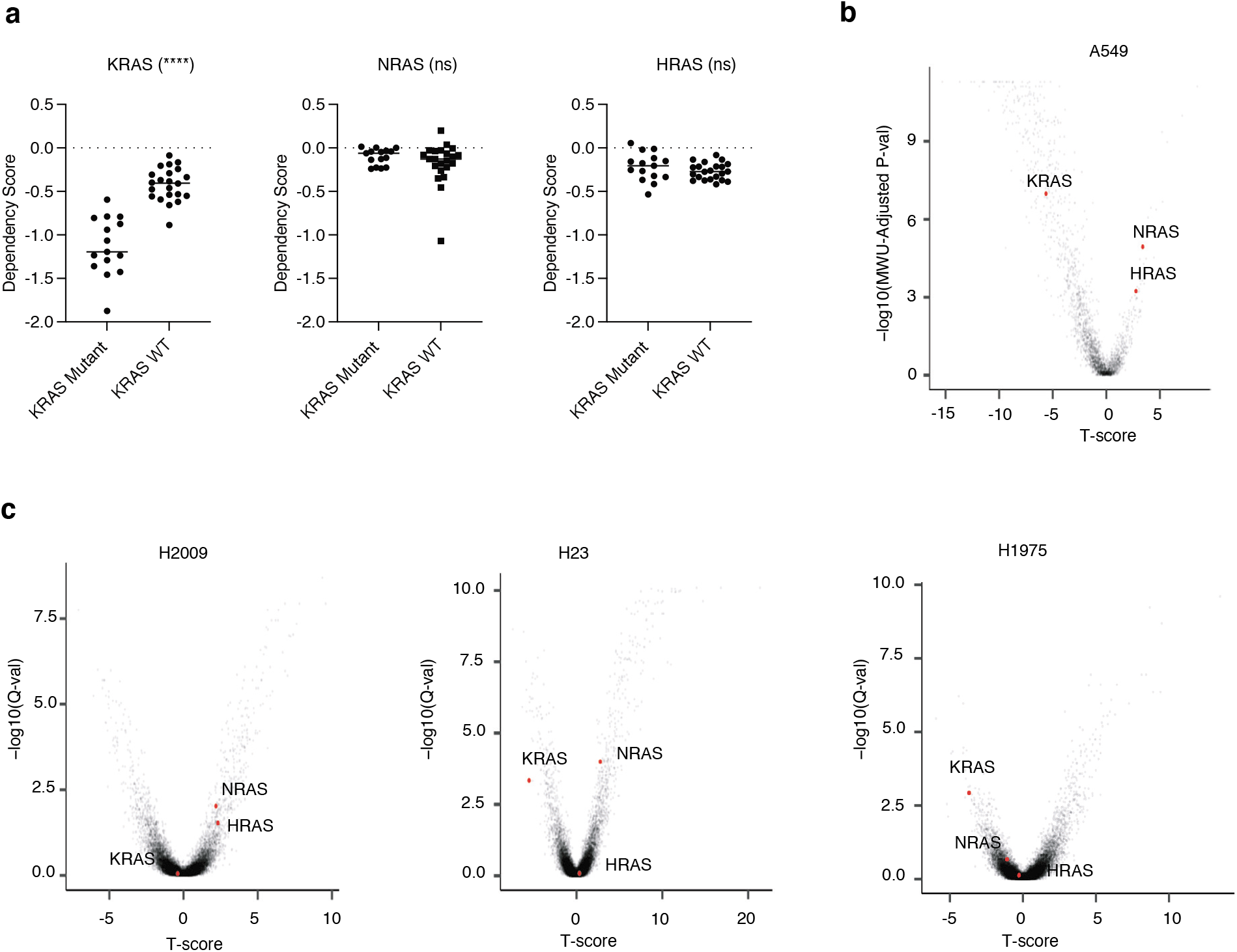
Dependency of human LUAD cell lines on RAS family members. **a.** Comparison of RAS family member dependency scores between KRAS mutant and KRAS wildtype human LUAD cell lines. **** (p <0.0001), ns (not-significant). **b.** Volcano plot showing the effects of RAS gene knockouts in A549 cells. The T-score represents the normalized effect of multiple sgRNAs targeting a gene. A positive T-score indicates a tumor suppressive effect. The effects of each gene relative to SAFE sgRNAs were tested via Mann–Whitney U (MWU) test, corrected via Benjamini-Hochberg procedure and shown as −log10(MWU-Adjusted P-val). (Data source: Kelly, Kostyrko, Han *et al.* 2020) **c.** Volcano plot showing effects of RAS gene knockouts in KRAS-mutant human LUAD cells (left: H2009, mid: H23, right: H1975) in 3D culture. The T-score represents the normalized effect of multiple sgRNAs targeting a gene. A positive T-score indicates a tumor suppressive effect. The effects of each gene relative to SAFE sgRNAs were tested via two-side t-test, corrected via Benjamini-Hochberg procedure and shown as −log10(Q-val). (Data source: Han *et al.* 2020)

**Supplemental Figure 6.**
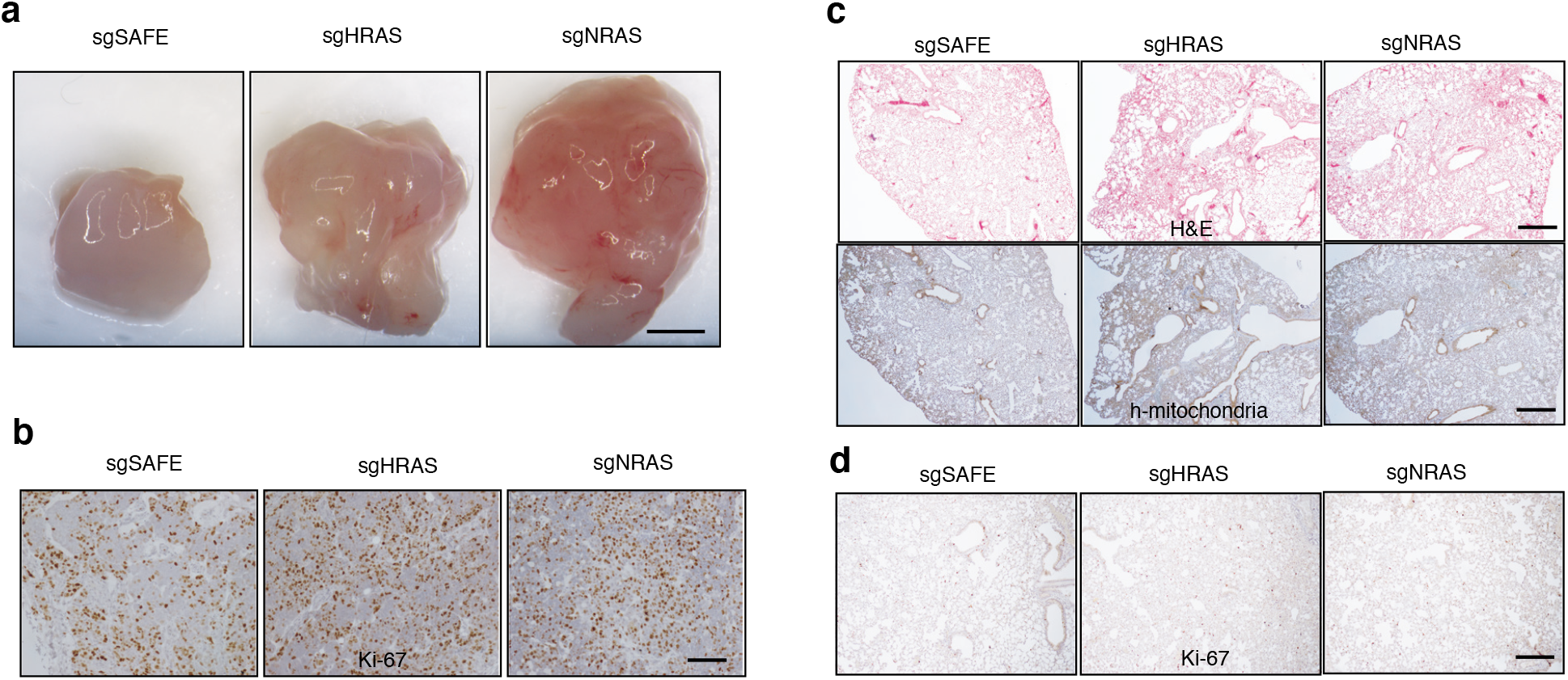
Inactivation of wild type HRAS or NRAS increases H23 cell growth after transplantation. **a.** Representative image of subcutaneous tumor size four weeks after transplantation with H23 cells as indicated. Quantification was shown in **Figure 3h**. Scale bar: 2 mm **b.** Representative image of Ki67 staining from subcutaneous tumor four weeks after transplantation with H23 cells as indicated. Quantification was shown in **Figure 3i**. Scale bar: 100 μm **c.** Representative image of HE (upper) and human mitochondria (lower) staining from lung tumor four weeks after intravenous transplantation with H23 cells as indicated. Quantification was shown in **Figure 3j**. Scale bar: 500 μm **d.** Representative image of Ki67 staining from lung tumor four weeks after intravenous transplantation with H23 cells as indicated. Quantification was shown in **Figure 3k**. Scale bar: 200 μm

**Supplemental Figure 7.**
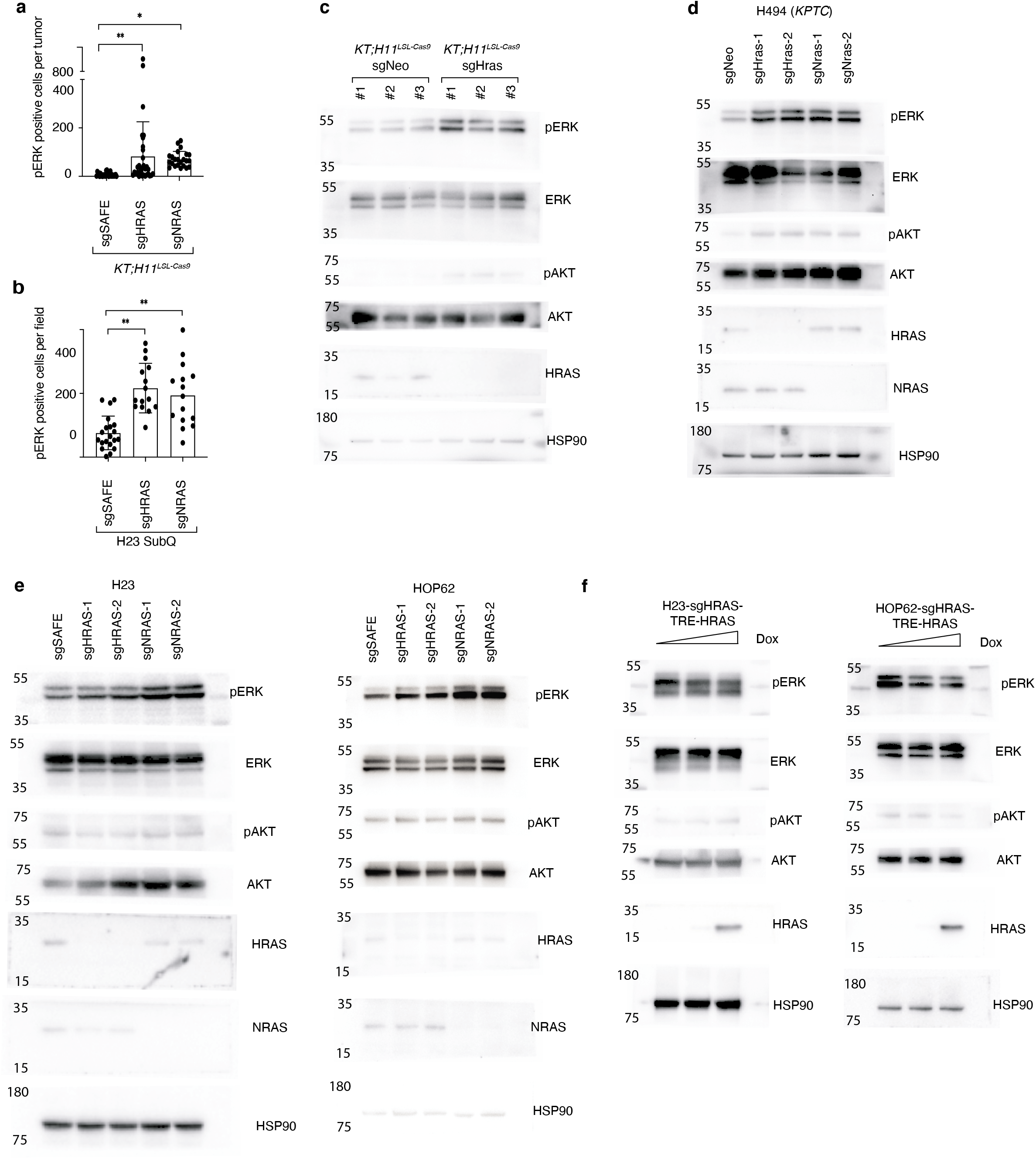
Wildtype RAS paralogs finetune RAS signaling. **a.** Quantification of pERK^pos^ cells in *KT;H11^LSL-Cas9^* mice with tumors initiated with Lenti-sgRNA/Cre vectors as indicated in **Figure 4a**. Each dot represents a tumor. *: p<0.05; **: p<0.01 **b.** Quantification of pERK^pos^ cells per field of indicated cells from **Figure 4b**. Each dot represents a view field. **: p<0.01 **c-f.** Raw images for western blots from **Figure 4c-f**. HRAS expression on **Figure 4f** were detected using same lysis on a different gel with increased loading.

**Supplemental Figure 8.**
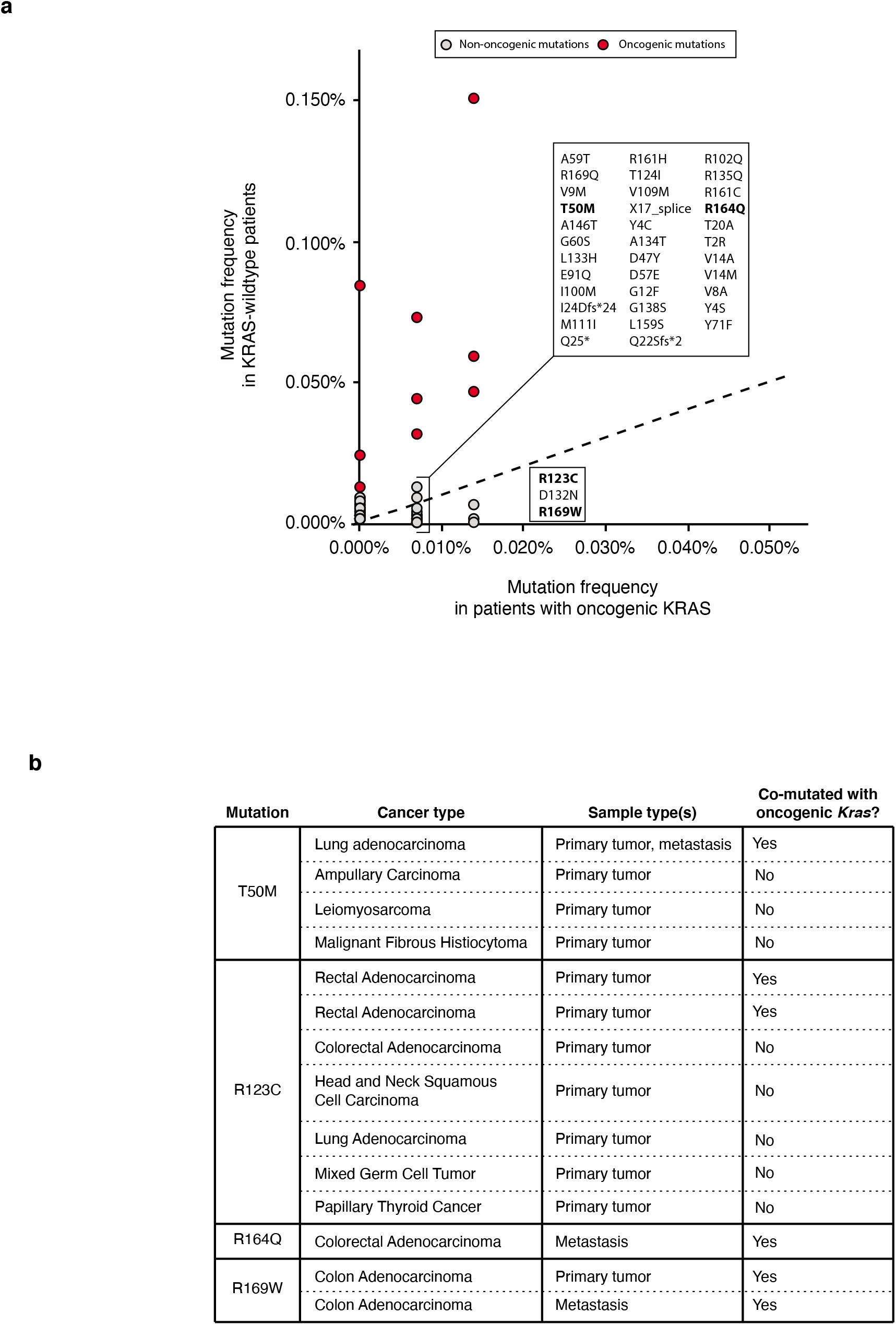
Identification of rare HRAS mutations in oncogenic KRAS-mutant tumors. **a.** Pan-cancer frequency of *HRAS* mutations in patients with *KRAS*-wildtype and oncogenic *KRAS*-mutant tumors from Project GENIE. Mutations that are intergenic, intronic, silent, or fall in the 3’ or 5’ UTR were excluded. Oncogenic *KRAS* mutants were defined as tumors having missense mutations in codons 12, 13 or 61. Known oncogenic *HRAS* mutations are highlighted in red. The dashed line indicates equal mutation frequency in *KRAS*-wildtype and mutant samples. Non-oncogenic mutations occurring at least once in patients with oncogenic *KRAS* mutations are annotated. HRAS mutants selected for analysis of ability to disrupt KRAS^G12D^-KRAS^G12D^ interactions are highlighted in bold. **b.** Characteristic of samples with rare HRAS mutants selected for analysis of their ability to disrupt KRAS^G12D^-KRAS^G12D^ interactions using the ReBiL2.0 system.

**Supplemental Figure 9.**
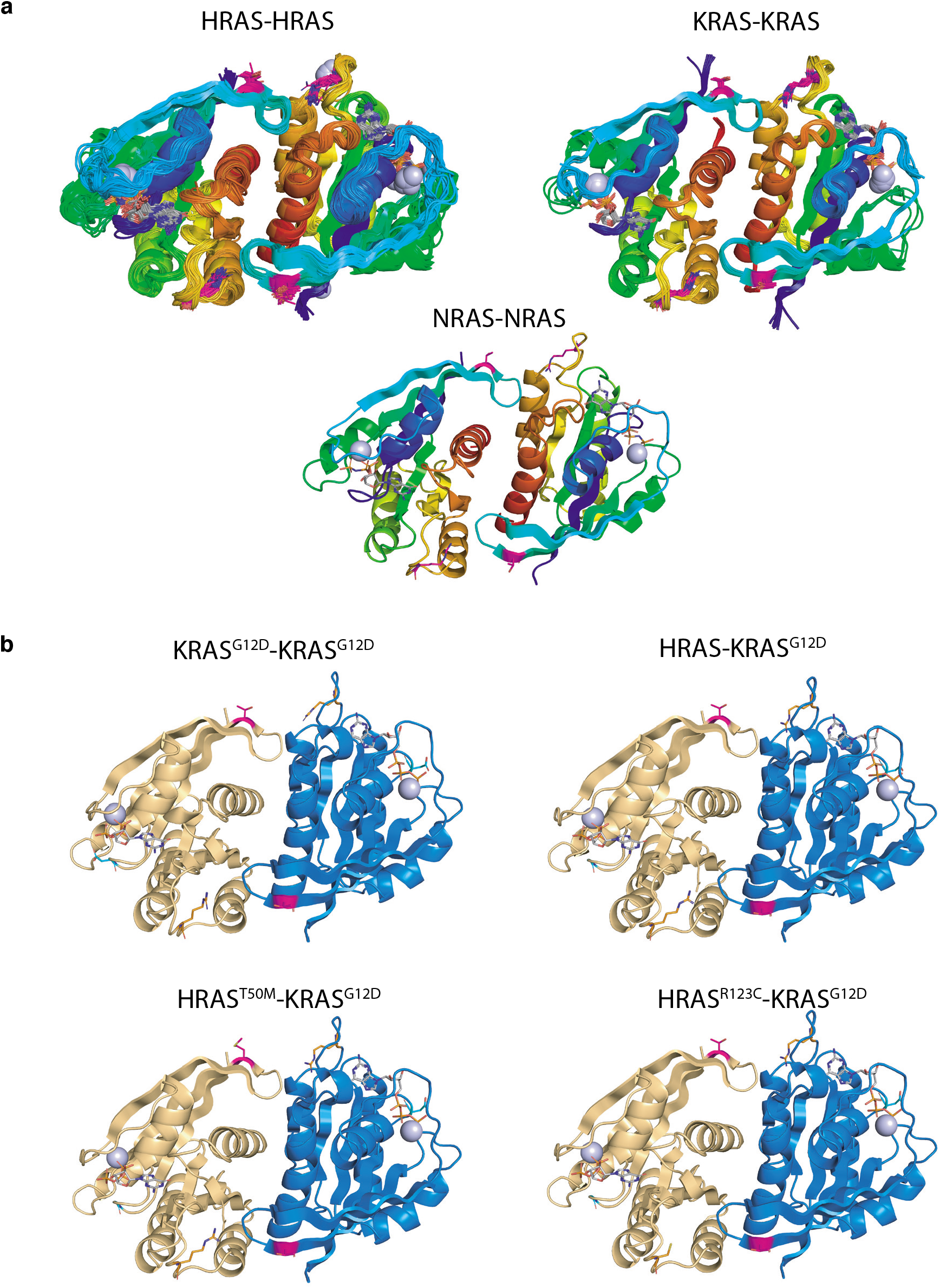
Modeling RAS-RAS dimer. **a.** Homodimers of RAS present in crystals of HRAS, KRAS, and NRAS in the Protein Data Bank. Dimers were downloaded from the Protein Common Interface Database (ProtCID)^58^, which clusters interfaces present in different crystals of homologous proteins. The *α*4-*α*5 dimer shown is present in 84 entries of HRAS, 13 entries of KRAS, and one entry of NRAS (PDB 5UHV). **b.** Models of a homodimer of KRAS^G12D^ and heterodimers of KRAS^G12D^ with HRAS, HRAS^T50M^, and HRAS^R123C^. The *α*4-*α*5 HRAS dimer from PDB entry 3K8Y was used as a template. KRAS^G12D^ from PDB entry 5USJ was superposed with the program PyMol on one or both monomers of 3K8Y to form the heterodimers and the homodimer respectively. Residues T50 and R123 were mutated with PyMol. All four structures were relaxed with the program Rosetta using the FastRelax protocol with the Ref2015 scoring function)^59^. Rosetta uses the backbone-dependent rotamer library of Shapovalov and Dunbrack to repack side chains around the mutated sites^60^. The resulting energies were: KRAS^G12D^-KRAS^G12D^, −1122.8 kcal/mol; HRAS-KRAS^G12D^, −1144.8 kcal/mol; HRAS^T50M^-KRAS^G12D^, −1135.5 kcal/mol; HRAS^R123C^-KRAS^G12D^, −1130.9 kcal/mol. Residues T50 (magenta) and R123 (orange) are indicated in sticks.

**Supplemental Figure 10.**
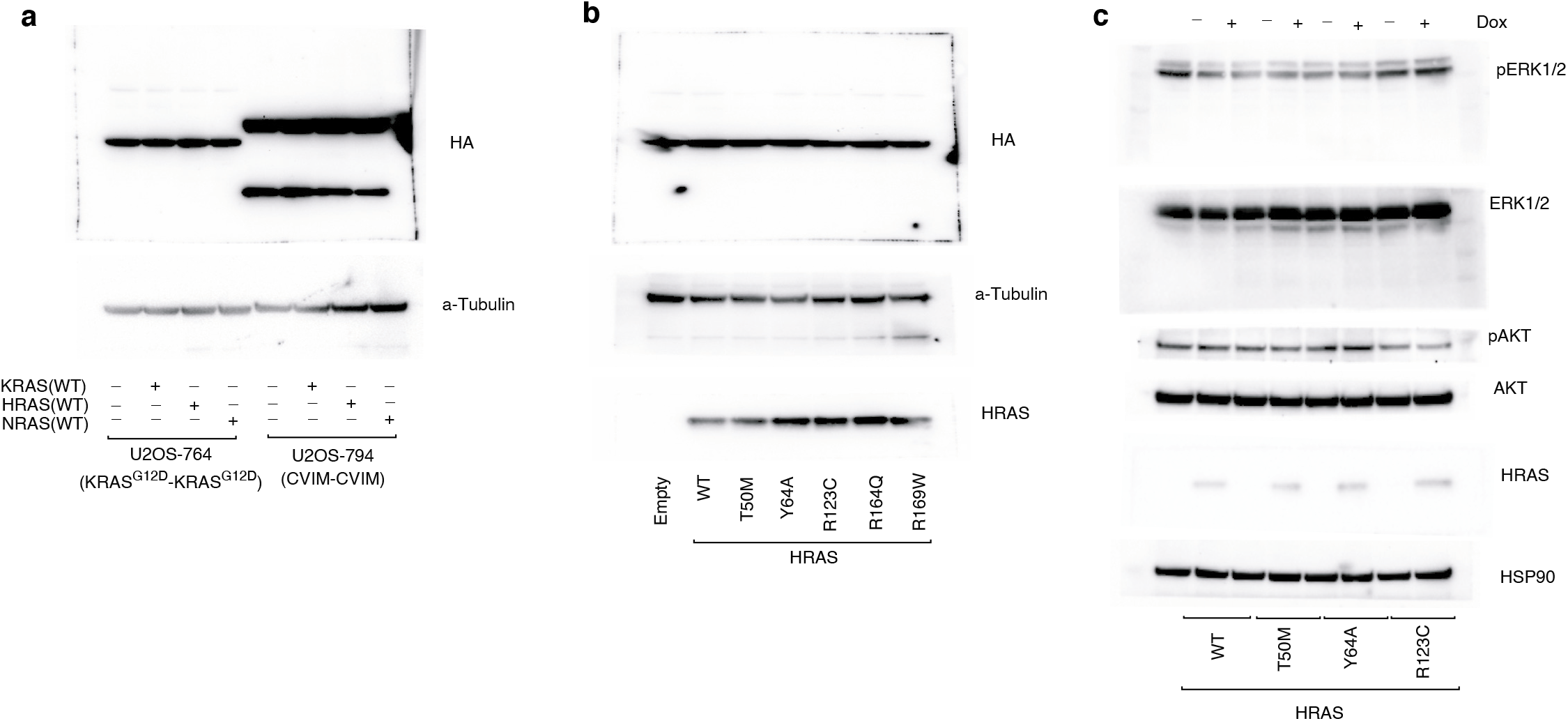
Wildtype RAS paralogs finetune RAS signaling through interaction with oncogenic KRAS. **a.** Raw images for western blots of split-luciferase (HA-tag) expression for ReBiL2.0 from **Figure 5c**. HA-tag expression were detected using same lysis on a different gel with increased loading. **b.** Raw images for western blots of split-luciferase (HA-tag) expression for ReBiL2.0 from **Figure 5e**. HA-tag expression were detected using same lysis on a different gel with increased loading. **c.** Raw images for western blots from **Figure 5h**. HRAS expression were detected using same lysis on a different gel with increased loading.

**Supplemental Figure 11.**
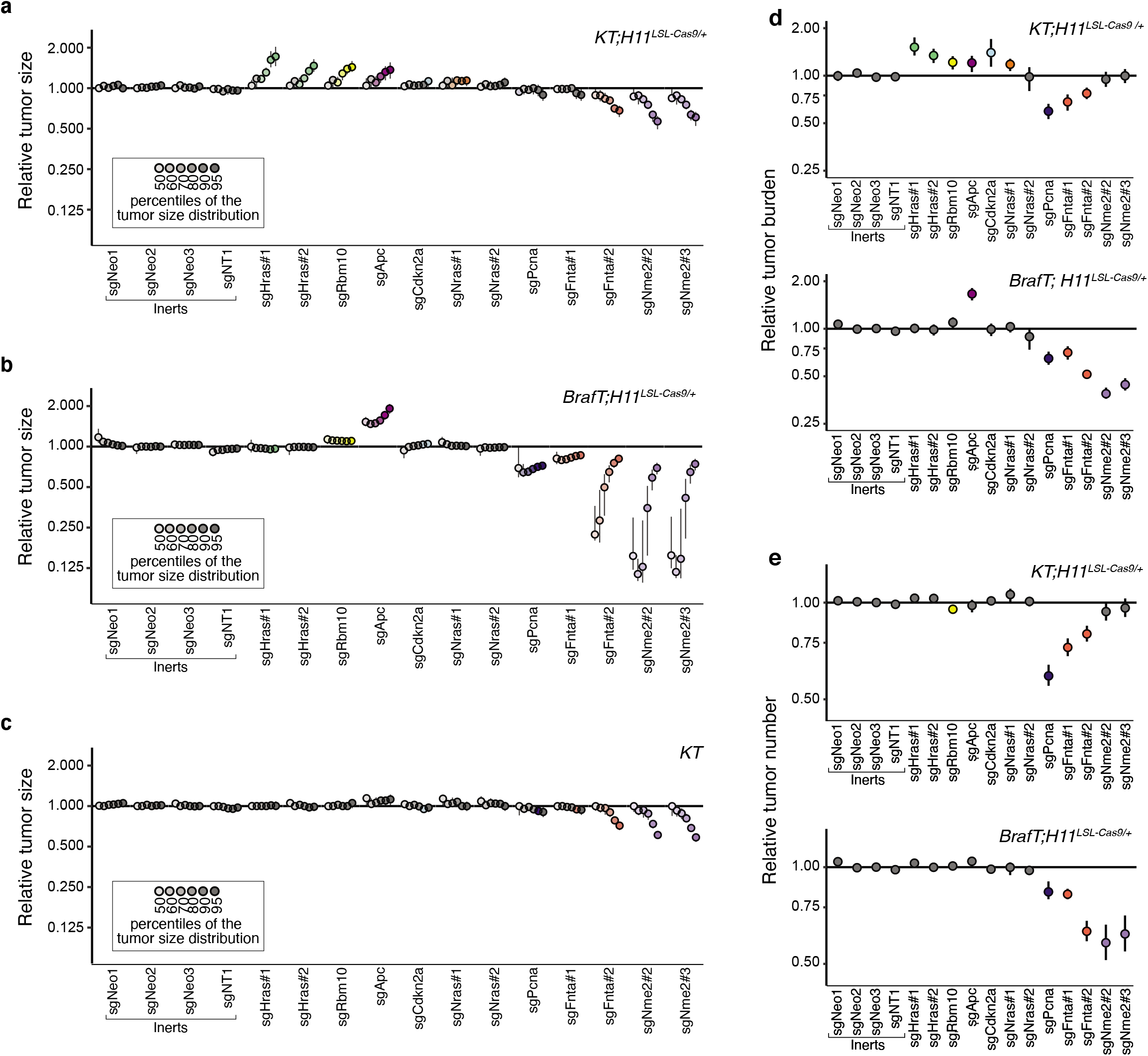
Paired screen in KRAS-driven and BRAF-driven lung cancer models validates HRAS and NRAS as KRAS-specific tumor suppressors. **a-c.** Tumor sizes at indicated percentiles for each sgRNA relative to the size of sgInert-containing tumors at the corresponding percentiles in *KT;H11^LSL-Cas9/+^* (**a**), BrafT;*H11^LSL-Cas9/+^* (**b**) and *KT* mice (**c**). Genes are ordered by 95^th^ percentile tumor size in *KT;H11^LSL-Cas9/+^* mice, with sgInerts on the left. sgInerts are in gray, and line at y=1 indicates no effect relative to sgInert. Error bars indicate 95% confidence intervals. Percentiles that are significantly different from sgInert (two-sided FDR-adjusted p < 0.05) are in color. Confidence intervals and P-values were calculated by bootstrap resampling. The negative effects of sgRNAs targeting *Fnta* and *Nme2* in the *KT* mice (**c**) are unexpected and indicate a potential bias in the size distributions of tumors with these genotypes. We note that the same bias may be present in the *KT;H11^LSL-Cas9/+^* and BrafT;*H11^LSL-Cas9/+^* data; however, sgRNAs targeting these genes in previous experiments showed consistent negative effects on tumor size, suggesting that the observed effects in this *KT;H11^LSL-Cas9/+^* cohort are not solely the product of this bias. **d.** The impact of each sgRNA on tumor burden relative to sgInerts in *KT;H11^LSL-Cas9/+^* (top) and BrafT;*H11^LSL-Cas9/+^* (bottom) mice, normalized to the corresponding statistic in *KT* mice to account for representation of each sgRNA in the viral pool. sgInerts are in gray and the line at y=1 indicates no effect. Error bars indicate 95% confidence intervals. Relative tumor burdens significantly different from sgInert (two-sided FDR-adjusted p<0.05) are in color. Confidence intervals and P-values were calculated by bootstrap resampling. **e.** The impact of each sgRNA on tumor number relative to sgInerts in *KT;H11^LSL-Cas9/+^* (top) and BrafT;*H11^LSL-Cas9/+^* (bottom) mice, normalized to the corresponding statistic in *KT* mice to account for representation of each sgRNA in the viral pool. sgInerts are in gray and the line at y=1 indicates no effect. Error bars indicate 95% confidence intervals. Relative tumor numbers significantly different from sgInert (two-sided FDR-adjusted p<0.05) are in color. Confidence intervals and P-values were calculated by bootstrap resampling.

## METHODS

### Cells, Reagents and Plasmids

H23, H727, and HOP62 cells were originally purchased from ATCC; HC494(*KPT*) lung adenocarcinoma cells were generated in the Winslow Lab; U2OS-134-764np (nl-KRAS^G12D^ cl-KRAS^G12D^; KRAS^G12D^ was fused to the N-termini of split luciferase proteins) and U2OS-134-794p (nl-CVIM cl-CVIM; CVIM represents the C-terminal last 20 amino acids of KRAS4B) cells were generated in the Wahl lab by Dr. Yao-Cheng Li (Salk Institute for Biological Studies). HC494 cells were cultured in DMEM containing 10% FBS, 100 units/mL penicillin and 100 μg/mL streptomycin. A549, H460 and H82 cells were cultured in RPMI1640 media containing 10% FBS, 100 units/mL penicillin and 100 μg/mL streptomycin. U2OS cells were cultured in DMEM/F12 (Thermo Fisher; phenol-red free), 10% (vol/vol) FBS, and 10 µg/mL ciprofloxacin. All cell lines were confirmed to be mycoplasma negative (MycoAlert Detection Kit, Lonza).

Trametinib was purchased from MedChemExpress (HY-10999); 5-Bromo-2′-deoxyuridine (10280879001) and D-Luciferin (L9504-5MG) was purchased from Sigma-Aldrich. All plasmids used in this study were listed in supplementary Table 1 and will be donated to Addgene.

### Design, generation, barcoding, and production of lentiviral vectors

The sgRNA sequences targeting the putative tumor suppressor genes were designed using CRISPick (https://portals.broadinstitute.org/gppx/crispick/public). All sgRNA sequence are shown in Supplementary Table 2. Each desired sgRNA vector was modified from our previously published pll3-U6-sgRNA-Pgk-Cre vector via site-directed mutagenesis (New England Biolabs, E0554S). The generation of the barcode fragment containing the 8-nucleotide sgID sequence and 20-nucleotide degenerate barcode, and subsequent ligation into the vectors were performed as previously described.

Lentiviral vectors were produced using polyethylenimine (PEI)-based transfection of 293T cells with delta8.2 and VSV-G packaging plasmids in 150-mm cell culture plates. Sodium butyrate (Sigma Aldrich, B5887) was added 8 hours after transfection to achieve a final concentration of 20 mM. Medium was refreshed 24 hours after transfection. 20 mL of virus-containing supernatant was collected 36, 48, and 60 hours after transfection. The three collections were then pooled and concentrated by ultracentrifugation (25,000 rpm for 1.5 hours), resuspended overnight in 100 µL PBS, then frozen at −80°C and were thawed and pooled at equal ratios immediately prior to delivery to mice.

### Mice and tumor initiation

The use of mice for the current study has been approved by Institutional Animal Care and Use Committee at Stanford University, protocol number 26696.

Kras^LSL-G12D/+^ (RRID:IMSR_JAX:008179), R26^LSL-tdTomato^ (RRID:IMSR_JAX:007909), and H11^LSL-Cas9^ (RRID:IMSR_JAX:027632) mice have been previously described. They were on a C57BL/6:129 mixed background. The B6.129P2(Cg)-Braf^tm1Mmcm^/J (BRAF^F-V600E^) mice were initially generated by Dankort et al. and obtained from the Jackson Laboratory (RRID:IMSR_JAX: 017837). Tumors were initiated by intratracheal delivery of 60 μl of lentiviral vectors dissolved in PBS.

For the initial experiments in Figure 1 and 2, tumors were allowed to develop for 12 weeks after viral delivery of a lentiviral pool that contained 19 barcoded Lenti-sgRNA/Cre vectors (Lenti-sgKrasIP/Cre). Tumors were initiated in Kras^LSL-G12D^; R26^LSL-tdTomato^ (*KT*), *KT;*H11^LSL-Cas9^; or *KT;p53^fl/fl^;*H11^LSL-Cas9^ mice with 1.95×10^5^ infectious units (ifu)/mouse.

For the validation experiments in Figure 3, tumors were allowed to develop for 15 weeks after viral delivery of a lentiviral pool that contained 26 barcoded Lenti-sgRNA/Cre vectors (Lenti-sgValidation/Cre). Tumors were initiated in Kras^LSL-G12D^; R26^LSL-tdTomato^ (*KT*) or *KT;*H11^LSL-Cas9^; mice with 3×10^5^ ifu/mouse.

For the individual sgRNA tumor initiation experiments in **Figure 3**, tumors were allowed to develop for 12 weeks after viral delivery of individual sgRNA expressing lentiviral vector that targeting Neo2, Hras, or Nras. Tumors were initiated in *KT;*H11^LSL-Cas9^; mice with 1×10^5^ ifu/mouse.

For the paired screen experiments in Figure 6, tumors were allowed to develop for 15 weeks after viral delivery of a lentiviral pool that contained 15 barcoded Lenti-sgRNA/Cre vectors (Lenti-sgMultiGEMM/Cre). Tumors were initiated in *KT;H11^LSL-Cas9/+^* or *Braf^V600E^;R26^LSL-tdTomato^;H11^LSL-Cas9/+^* mice with 3×10^5^ ifu/mouse. Note that *KT;*H11^LSL-Cas9/+^ rather than *KT;*H11^LSL-Cas9/LSL-Cas9^ mice were used in this experiment to match the Cas9 dosage of the *BrafT;H11^LSL-Cas9/+^* mice, whereas *KT;*H11^LSL-Cas9/LSL-Cas9^ mice were used in all other experiments. To evaluate the effects of Cas9 dosage on the tumor suppressive effects of the Lenti-sgMultiGEMM/Cre pool, we also initiated tumors in a small cohort of *KT;H11^LSL-Cas9/LSL-Cas9^* mice. Reductions in the magnitude of the effects of various sgRNAs were observed in the *KT;H11^LSL-Cas9/+^* cohort relative to the *KT;H11^LSL-Cas9/LSL-Cas9^* cohort, underscoring the importance of matching Cas9 dosage and suggesting that Cas9 can be limiting in H11LSL-Cas9/+ mice.

### Tuba-seq library generation

Genomic DNA was isolated from bulk tumor-bearing lung tissue from each mouse as previously described. Briefly, benchmark control cell lines were generated from LSL-YFP MEFs transduced by a barcoded Lenti-sgNT3/Cre vector (NT3: an inert sgRNA with a distinct sgID) and purified by sorting YFP^pos^ cells. Three benchmark control cell lines (500,000 cells each) were added to each mouse lung sample prior to lysis to enable the calculation of the absolute number of neoplastic cells in each tumor from the number of sgID-BC reads. Following homogenization and overnight protease K digestion, genomic DNA was extracted from the lung lysates using standard phenol-chloroform and ethanol precipitation methods. Subsequently, Q5 High-Fidelity 2x Master Mix (New England Biolabs, M0494X) was used to amplify the sgID-BC region from 32 μg of genomic DNA in a total reaction volume of 800 μl per sample. The unique dual-indexed primers used were Forward: AAT GAT ACG GCG ACC ACC GAG ATC TAC AC-8 nucleotides for i5 index-ACA CTC TTT CCC TAC ACG ACG CTC TTC CGA TCT-6 to 9 random nucleotides for increased diversity-GCG CAC GTC TGC CGC GCT G and Reverse: CAA GCA GAA GAC GGC ATA CGA GAT-6 nucleotides for i7 index-GTG ACT GGA GTT CAG ACG TGT GCT CTT CCG ATC T-9 to 6 random nucleotides for increased diversity-CAG GTT CTT GCG AAC CTC AT. The PCR products were purified with Agencourt AMPure XP beads (Beckman Coulter, A63881) using a double size selection protocol. The concentration and quality of the purified libraries were determined using the Agilent High Sensitivity DNA kit (Agilent Technologies, 5067-4626) on the Agilent 2100 Bioanalyzer (Agilent Technologies, G2939BA). The libraries were pooled based on lung weight to ensure even reading depth, cleaned up again using AMPure XP beads, and sequenced (read length 2×150bp) on the Illumina HiSeq 2500 or NextSeq 550 platform (Admera Health Biopharma Services).

### Generation of Stable Cell Lines

Parental cells were seeded at 50% confluency in a 6-well plate the day before transduction (day 0). The cell culture medium was replaced with 2 mL fresh medium containing 8 µg/mL hexadimethrine bromide (Sigma Aldrich, H9268-5G), 20 µL ViralPlus Transduction Enhancer (Applied Biological Materials Inc., G698) and 40 µL concentrated lentivirus and cultured overnight (Day 1). The medium was then replaced with complete medium and cultured for another 24 hours (Day 2). Cells were transferred into a 100 mm cell culture dish with appropriate amounts of antibiotic (Blasticidin doses: U2OS: 10 µg/mL; HOP62: 50 µg/mL; H727: 10 µg/mL; H23: 15 µg/mL; Puromycin doses: HC494: 5 µg/mL; U2OS: 1 µg/mL; HOP62: 5 µg/mL; H727: 5 µg/mL; H23: 5 µg/mL) and selected for 48 hours (Day 3).

### Western Blot

Cells were lysed in RIPA buffer (50 mM Tris-HCl (pH 7.4), 150 mM NaCl, 1% Nonidet P-40, and 0.1% SDS) and incubated at 4 °C with continuous rotation for 30 minutes, followed by centrifugation at 12,000 × rcf for 10 minutes. The supernatant was collected, and the protein concentration was determined by BCA assay (Thermo Fisher Scientific, 23250). Protein extracts (10–50 μg) were dissolved in 10% SDS-PAGE and transferred onto PVDF membranes. The membranes were blocked with 5% non-fat milk in TBS with 0.1% Tween 20 (TBST) at room temperature for one hour, cut according to the molecular weight of target protein (with at least two flacking protein marker), followed by incubation with primary antibodies diluted in TBST (1:1000) at 4 °C overnight. After three 10-minutes washes with TBST, the membranes were incubated with the appropriate secondary antibody conjugated to HRP diluted in TBST (1:10000) at room temperature for 1 hour. After three 10-minutes washes with TBST, Protein expression was quantified with enhanced chemiluminescence reagents (Fisher Scientific, PI80196). For AKT and ERK, phosphorylated proteins were detected first and the membrane were striped, blocked, and incubated with 1^st^ and 2^nd^ antibodies for pan protein detections.

Antibodies used in this study: HSP90 (BD Biosciences, 610418), pAKT (Cell Signaling, 4060S), pERK (Cell Signaling, 4370L), ERK (Cell Signaling, 9102S), AKT (Cell Signaling, 4691S), HRAS (Thermo Fisher Scientific, 18295-1-AP), NRAS (Santa Cruz Biotechnology, sc-31), HA-tag (Santa Cruz Biotechnology, sc-7392).

### Histology and immunohistochemistry (IHC)

Lung lobes were fixed in 4% formalin and paraffin embedded. Hematoxylin and eosin staining was performed using standard methods. IHC was performed on 4-μm sections with IHC was performed using Avidin/Biotin Blocking Kit (Vector Laboratories, SP-2001), Avidin-Biotin Complex kit (Vector Laboratories, PK-4001), and DAB Peroxidase Substrate Kit (Vector Laboratories, SK-4100) following standard protocols.

The following primary antibodies were used: Ki-67 (BD Pharmingen, 550609), BrdU (BD Pharmingen, 555627), human mitochondria (Abcam, ab92824), pERK (Cell Signaling, 4370L). Total tumor burden (tumor area/total area × 100%), mitochondria^pos^ tumor burden (mitochondria^pos^ area/total area × 100%), BrdU^pos^ cell number, Ki67^pos^ cell number, and pERK^pos^ cell number were calculated using ImageJ.

### Cell proliferation assay (CCK8)

For cell proliferation assays, cells were seeded in 96-well plates at a density of 5000 cells per well and allowed to adhere overnight in regular growth media (Day 0). Cells were then cultured in media as indicated on each figure panel for 7 days. Relative cell number were measured every other day using Cell Counting Kit-8 (Bimake, B34304) according to the manufacturer’s instructions.

### Colony formation assay

For clonogenic assays, cells were seeded in 6-well plates at a density of 500 cells per well and allowed to adhere overnight in regular growth media. Cells were then cultured in media as indicated on each figure panel for 14 days. Growth media with or without drugs was replaced every 2 days. At the end point, cells were stained with 0.5% crystal violet in 20% methanol. Colony numbers were calculated using ImageJ

### Allograft studies in immunocompromised mice

For intravenous transplants into immunocompromised NSG mice, 5×10^5^ H23 cells were injected into one of lateral tail veins. Mice were sacrificed 28 days post-injection and lung lobe were fixed in 4% formalin and paraffin embedded. For subcutaneous transplants into immunocompromised NSG mice, 2× 10^6^ of each H23 cells (sgSAFE, sgHRAS, and sgNRAS) were re-suspended in 200uL Matrigel^®^ Basement Membrane Matrix (Corning, 354234) and injected into three parallel sites per mouse. Mice were sacrificed 28 days post-injection. Tumors were dissected and the weight, height, width, and length, of each tumor was measured. Tumor volume was roughly calculated via the formula: V = (4/3) × π × (L/2) × (L/2) × (D/2).

Institute of Medicine Animal Care and Use Committee approved all animal studies and procedures.

### ReBiL2.0 assay

ReBiL2.0 assay was performed as previously descried^16^. ReBiL cells (U2OS-134-764np or U2OS-134-794p with overexpression of KRAS4b, HRAS or NRAS) were seeded in i) 96-well plates at density of 2×10^4^, and ii) 6-well plates at density of 1×10^6^ and allowed to adhere overnight in regular growth media (DMEM/F12, 10% FBS, and 10 µg/mL ciprofloxacin). The next day, cells were then cultured in serum limited media (DMEM/F12, 1% FBS, and 10 µg/mL ciprofloxacin) containing 100 ng/mL doxycycline for 24 hours. Upon termination of the ReBiL assay, i) to measure raw luciferase activity, 300 µM D-luciferin was added to 96-well plate culture and incubate in 37°C for 30mins and raw luminescent data collected by a Tecan microplate reader; ii) to measure viable cell numbers, CCK-8 assay were performed in the same 96-well plate culture and raw cell number data collected by a Tecan microplate reader; iii) to quantify the 1/2luc fusion proteins, ReBiL cells from 6-well plate culture were harvested with RIPA lysis buffer for protein extraction and western blot was performed for HA-tag and HSP90 expression. Then the ReBiL2.0 score was calculated via the formula:

ReBiL2.0 score = ([Raw Luminescence]/[Cell number]) / ([1/2luc Least]/[HSP90])

### Analysis of human lung adenocarcinoma cancer genome sequencing data (for HRAS rare mutations)

To assess evidence that *HRAS* functions as a Kras-specific tumor suppressor in human cancer, we queried publicly available cancer genomic datasets. GENIE Release 9.1-public was accessed through the Synapse platform and data on somatic mutations (data_mutations_extended.txt), sample- and patient-level clinical data (data_clinical_sample.txt and data_clinical_patient.txt), and genotyping panel information (genomic_information.txt) were downloaded. While it is unclear how our findings may extrapolate to cancer types beyond lung adenocarcinoma, Hras mutations are exceedingly rare (occurring at a frequency of just ∼0.008 in GENIE samples) so we performed a pan-cancer analysis. Each sample was assigned to its patient of origin and annotated for the presence of both oncogenic Kras mutations (defined as missense mutations in Kras exons 12, 13 or 61) and for the presence of potentially functional Hras mutations (variants that were silent, intergenic, intronic, or fell in the 3’ or 5’ UTRs were excluded from this analysis). When multiple samples were derived from the same patient, the patient in question was annotated as having a mutation if it occurred in at least one of their associated samples. From this information we produced a list of the frequency of all Hras variants in patients with and without oncogenic Kras in both datasets. The genotyping panel information was used to identify GENIE patients that were not genotyped at Hras and exclude these from the frequency calculation.

### Process paired-end reads to identify the sgID and barcode

Sequencing of Tuba-seq libraries produces reads that are expected to contain an 8-nucleotide sgID followed by a 30-nucleotide barcode (BC) of the form GCNNNNNTANNNNNGCNNNNNTANNNNNGC, where each of the 20 Ns represent random nucleotides. Each sgID has a one-to-one correspondence with an sgRNA in the viral pool; thus, the sgID sequence identifies the gene targeted in a given tumor. Note that all sgID sequences in the viral pool differ from each other by at least three nucleotides such that incorrect sgID assignment (and thus, inference of tumor genotype) due to PCR or sequencing error is extremely unlikely. The random 20-nucleotide portion of the BC is expected to be unique to each lentiviral integration event, and thus tags all cells in a single clonal expansion. Note that the length of the barcode ensures a high theoretical potential diversity (∼4^20^ > 10^12^ barcodes per vector), so while the actual diversity of each Lenti-sgRNA/Cre vector is dictated by the number of colonies generated during the plasmid barcoding step, it is very unlikely that we will observe the same BC in multiple clonal expansions.

FASTQ files were parsed using regular expressions to identify the sgID and BC for each read. To minimize the effects of sequencing error on BC identification, we required the forward and reverse reads to agree completely within the 30-nucleotide sequence to be further processed. We also screened for barcodes that were likely to have arisen due to errors in sequencing the barcodes of genuine tumors. Given the low rate of sequencing error, we expect these spurious “tumors” to have read counts that are far lower than the read counts of the genuine tumors from which they arise. While it is impossible to eliminate these spurious tumors, we sought to minimize their effect by identifying small “tumors” with barcodes that are highly similar to the barcodes of larger tumors. Specifically, if a pair of “tumors” had barcodes that were within a Hamming distance of two, and if one of the tumors had less than 5% as many reads as the other, then the reads associated with the smaller tumor were attributed to the larger tumor.

After these filtering steps, the read counts associated with each barcode were converted to absolute neoplastic cell numbers by normalizing to the number of reads in the “spike-in” cell lines added to each sample prior to lung lysis and DNA extraction. The median sequencing depth across experiments was ∼1 read per 6.4 cells.

For statistical comparisons of tumor genotypes, we applied a minimum tumor size cutoff of 100 cells. In selecting a cutoff, we sought to include tumors that are large enough to be consistently detected despite differences in sequencing depth among mice, while using as many tumors as possible to maximize the statistical power. Importantly, we analyzed each Tuba-seq dataset with multiple minimum tumor size cut-offs (50, 100, 200, 500 cells) and found that our findings were robust.

### Summary statistics for overall growth rate

To assess the extent to which a given gene (*X)* affects tumor growth, we compared the distribution of tumor sizes produced by vectors targeting that gene (sg*X* tumors) to the distribution produced by our negative control vectors (sg*Inert* tumors). We relied on two statistics to characterize these distributions: the size of tumors at defined percentiles of the distribution (specifically the 50^th^, 60^th^, 70^th^, 80^th^, 90^th^, and 95^th^ percentile tumor sizes), and the log-normal mean size (LN mean). The percentile sizes are nonparametric summary statistics of the tumor size distribution. In considering percentiles corresponding to the right tail of the distribution, we focus on the growth of larger tumors, thereby avoiding issues stemming from potential variation in cutting efficiency among guides. The LN mean is the maximum-likelihood estimate of mean tumor size assuming a log-normal distribution. Previous work found that this statistic represents the best parametric summary of tumor growth based on the maximum likelihood quality of fit of various common parametric distributions.

To quantify the extent to which each gene suppressed or promoted tumor growth, we normalized statistics calculated on tumors of each genotype to the corresponding inert statistic. The resulting ratios reflect the growth advantage (or disadvantage) associated with each tumor genotype relative to the growth of *sgInert* tumors.

For example, the relative i^th^ percentile size for tumors of genotype X was calculated as:

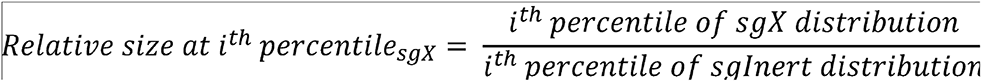

Likewise, the relative LN mean size for tumors of genotype X was calculated as:

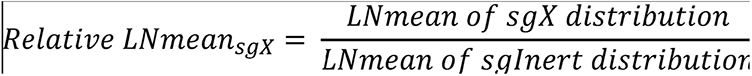

### Summary statistics for relative tumor number and relative tumor burden

In addition to the tumor size metrics described above, we characterized the effects of gene inactivation on tumorigenesis in terms of the number of tumors and total neoplastic cell number (“tumor burden”) associated with each genotype. Unlike the aforementioned metrics of tumor size, tumor number and burden are linearly affected by lentiviral titer and are thus sensitive to underlying differences in the representation of each Lenti-sgRNA/Cre vector in the viral pool. Critically, each Tuba-seq experiment included a cohort of *KT* control mice. *KT* mice lack expression of Cas9, thus all Lenti-sgRNA/Cre vectors are functionally equivalent in these mice, and the observed tumor number and burden associated with each sgRNA reflects the make-up of the viral pool.

To assess the extent to which a given gene (*X)* affects tumor number, we therefore first normalized the number of sgX tumors in *KT;H11^LSL-Cas9^* mice (also *KT;p53^flox/flox^;H11^LSL-Cas9^* and *Braf^LSL-V600E/+^T; H11^LSL-Cas9^* mice in the initial Kras-interacting protein screen and the paired screen, respectively) to the number of sgX tumors in the *KT* mice:

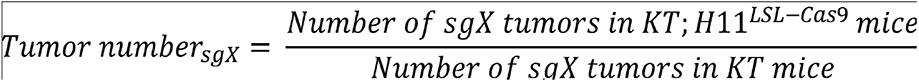

As with the tumor size metrics, we then calculated a relative tumor number by normalizing this statistic to the corresponding statistic calculated using sgInert tumors:

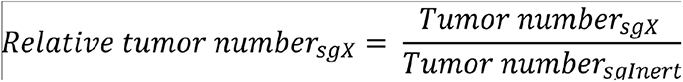

Genes that influence relative tumor number modify the probability of tumor initiation and/or the very early stages of oncogene-driven epithelial expansion, which prior work suggests are imperfectly correlated with tumor growth at later stages. Relative tumor number thus captures an additional and potentially important aspect of tumor suppressor gene function.

Analogous to the calculation of relative tumor number, we characterized the effect of each gene on tumor burden by first normalizing the sgX tumor burden in Cas9-expressing mice to the burden in KT mice:

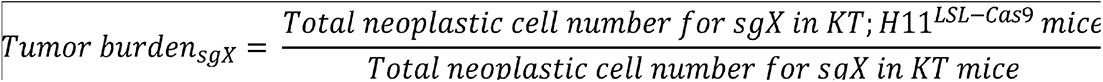

We then calculated a relative tumor burden by normalizing this number to the corresponding statistic calculated using sgInert tumors:

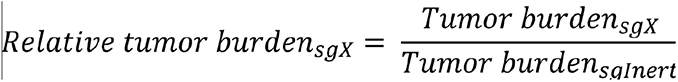

Tumor burden is an integration over tumor size and number, and thus reflects the total neoplastic load in each mouse. Tumor burden is thus more strongly related to morbidity than are our metrics of tumor size and is closely related to traditional measurements of tumor progression such as duration of survival and tumor area. While intuitively appealing, tumor burden is notably nosier than our metrics of tumor size as it is strongly determined by the size of the largest tumors.

### Calculation of confidence intervals and P-values for tumor growth and number metrics

Confidence intervals and *P*-values were calculated using bootstrap resampling to estimate the sampling distribution of each statistic. To account for both mouse-to-mouse variability and variability in tumor size and number within mice, we adopted a two-step, nested bootstrap approach where we first resampled mice, and then resampled tumors within each mouse in the pseudo-dataset. 10,000 bootstrap samples were drawn for all reported P-values. 95% confidence intervals were calculated using the 2.5^th^ and 97.5^th^ percentile of the bootstrapped statistics. Because we calculate metrics of tumor growth that are normalized to the same metrics in sgInert tumors, under the null model where genotype does not affect tumor growth, the test statistic is equal to 1. Two-sided p-values were thus calculated as followed:

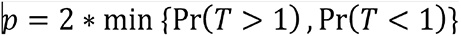

Where T is the test statistic and Pr(T>1) and Pr(T<1) were calculated empirically as the proportion of bootstrapped statistics that were more extreme than the baseline of 1. To account for multiple hypothesis testing, p-values were FDR-adjusted using the Benjamini-Hochberg procedure as implemented in the Python package stats models.

### AP-MS data visualization

AP-MS data was analyzed as described (Ding et al 2016). Briefly, protein spectral matches (PSMs; Kelly et al 2020) were normalized by protein length and total spectral matches per experiment. These normalized spectral abundance factors (NSAFs) were then normalized to NSAFs of matched prey proteins from a large cohort of unrelated AP/MS experiments to produce a Z-score. Z-scores are proportional to the areas of circles in bubble plots. In cluster diagrams, NSAFs are binarized by statistical significance (FDR > 0.5), similarities between interactome profiles are determined by cosine distance, and dendrogram topology is determined by UPGMA.

### Modeling RAS-RAS dimer

Potential templates for modeling the heterodimers were obtained from the ProtCID database. ProtCID is built from clustering interfaces of homologous proteins obtained from domain-domain contacts within protein crystals in the Protein Data Bank. Hierarchical clustering of interfaces is performed with a Jaccard-index similarity metric based on the contacts shared between different interfaces. Models for the structure of the HRAS/KRAS heterodimer were built by superposing a structure of KRAS-G12D (PDB: 5USJ) onto a monomer of the HRAS homodimer in PDB entry 3K8Y.

## ACKNOWLEDGEMNTS

We thank the Stanford Veterinary Animal Care Staff for expert animal care, Human Pathology/Histology Service Center, Stanford Protein and Nucleic Acid Facility for experimental support; A. Orantes for administrative support; Members of the Winslow laboratory and Ian Prior for helpful comments. R.T. was supported by a Stanford University School of Medicine Dean’s Postdoctoral Fellowship, a Tobacco-Related Disease Research Program (TRDRP) Postdoctoral fellowship (27FT-0044), and a Stanford Cancer Institute fellowship. C.W.M. was supported by the NSF Graduate Research Fellowship Program and an Anne T. and Robert M. Bass Stanford Graduate Fellowship. J.D.H was supported by a Stanford University School of Medicine Dean’s Postdoctoral Fellowship and a TRDRP Postdoctoral fellowship (T31FT1619). H.C. was supported by a TRDRP Postdoctoral Fellowship (28FT-0019). N.W.H. was supported by the NSF Graduate Research Fellowship Program. Work in the laboratory of G.M.W. was supported, in part, by Cancer Center Core Grant CA014195, the Breast Cancer Research Foundation, the Freeberg Foundation and the NIH/National Cancer Institute (Grant R35 CA197687). This work was supported by NIH R01-CA230025 (to M.M.W), NIH R01-CA231253 (to M.M.W and D.A.P), NIH R01-CA234349 (to M.M.W and D.A.P.), TRDRP 27IP-0052 (to M.M.W), and in part by the Stanford Cancer institute support grant (NIH P30-CA124435).

## REFERENCES

1. Karnoub, A.E. & Weinberg, R.A. Ras oncogenes: split personalities. Nature reviews Molecular cell biology 9, 517–531 (2008).

2. Cox, A.D., Fesik, S.W., Kimmelman, A.C., Luo, J. & Der, C.J. Drugging the undruggable RAS: mission possible? Nature reviews Drug discovery 13, 828–851 (2014).

3. Zhou, B., Der, C.J. & Cox, A.D. in Seminars in cell & developmental biology, Vol. 58 60–69 (Elsevier, 2016).

4. Wennerberg, K., Rossman, K.L. & Der, C.J. The Ras superfamily at a glance. Journal of cell science 118, 843–846 (2005).

5. Hobbs, G.A., Der, C.J. & Rossman, K.L. RAS isoforms and mutations in cancer at a glance. Journal of cell science 129, 1287–1292 (2016).

6. Stephen, A.G., Esposito, D., Bagni, R.K. & McCormick, F. Dragging ras back in the ring. Cancer cell 25, 272–281 (2014).

7. Brose, M.S. et al. BRAF and RAS mutations in human lung cancer and melanoma. Cancer research 62, 6997–7000 (2002).

8. Prior, I.A., Lewis, P.D. & Mattos, C. A comprehensive survey of Ras mutations in cancer. Cancer research 72, 2457–2467 (2012).

9. Papke, B. & Der, C.J. Drugging RAS: Know the enemy. Science 355, 1158–1163 (2017).

10. Kelly, M.R. et al. Combined proteomic and genetic interaction mapping reveals new RAS effector pathways and susceptibilities. Cancer discovery 10, 1950–1967 (2020).

11. Broyde, J. et al. Oncoprotein-specific molecular interaction maps (SigMaps) for cancer network analyses. Nature biotechnology 39, 215–224 (2021).

12. Zhou, Y. & Hancock, J.F. Deciphering lipid codes: K-Ras as a paradigm. Traffic 19, 157–165 (2018).

13. Wittinghofer, A. & Pal, E.F. The structure of Ras protein: a model for a universal molecular switch. Trends in biochemical sciences 16, 382–387 (1991).

14. Omerovic, J., Hammond, D.E., Clague, M.J. & Prior, I.A. Ras isoform abundance and signalling in human cancer cell lines. Oncogene 27, 2754–2762 (2008).

15. Han, K. et al. CRISPR screens in cancer spheroids identify 3D growth-specific vulnerabilities. Nature 580, 136–141 (2020).

16. Li, Y.-C. et al. Analysis of RAS protein interactions in living cells reveals a mechanism for pan-RAS depletion by membrane-targeted RAS binders. Proceedings of the National Academy of Sciences 117, 12121–12130 (2020).

17. Hingorani, S.R. et al. Preinvasive and invasive ductal pancreatic cancer and its early detection in the mouse. Cancer cell 4, 437–450 (2003).

18. Cai, H. et al. A functional taxonomy of tumor suppression in oncogenic KRAS-driven lung cancer. Cancer Discovery (2021).

19. Rogers, Z.N. et al. Mapping the in vivo fitness landscape of lung adenocarcinoma tumor suppression in mice. Nature genetics 50, 483–486 (2018).

20. Rogers, Z.N. et al. A quantitative and multiplexed approach to uncover the fitness landscape of tumor suppression in vivo. Nature methods 14, 737–742 (2017).

21. Chuang, C.-H. et al. Molecular definition of a metastatic lung cancer state reveals a targetable CD109–Janus kinase–Stat axis. Nature medicine 23, 291–300 (2017).

22. Ruiz, S., Santos, E. & Bustelo, X.R. RasGRF2, a guanosine nucleotide exchange factor for Ras GTPases, participates in T-cell signaling responses. Molecular and cellular biology 27, 8127–8142 (2007).

23. Brandt, A.C., Koehn, O.J. & Williams, C.L. SmgGDS: An Emerging Master Regulator of Prenylation and Trafficking by Small GTPases in the Ras and Rho Families. Frontiers in Molecular Biosciences 8, 542 (2021).

24. Rowell, C.A., Kowalczyk, J.J., Lewis, M.D. & Garcia, A.M. Direct demonstration of geranylgeranylation and farnesylation of Ki-Ras in vivo. Journal of Biological Chemistry 272, 14093–14097 (1997).

25. Zhang, F.L. et al. Characterization of Ha-ras, N-ras, Ki-Ras4A, and Ki-Ras4B as in vitro substrates for farnesyl protein transferase and geranylgeranyl protein transferase type I. Journal of Biological Chemistry 272, 10232–10239 (1997).

26. Takaya, A. et al. R-Ras regulates exocytosis by Rgl2/Rlf-mediated activation of RalA on endosomes. Molecular biology of the cell 18, 1850–1860 (2007).

27. Marais, R., Light, Y., Paterson, H. & Marshall, C. Ras recruits Raf-1 to the plasma membrane for activation by tyrosine phosphorylation. The EMBO journal 14, 3136–3145 (1995).

28. Campbell, J.D. et al. Distinct patterns of somatic genome alterations in lung adenocarcinomas and squamous cell carcinomas. Nature genetics 48, 607–616 (2016).

29. Sánchez-Rivera, F.J. et al. Rapid modelling of cooperating genetic events in cancer through somatic genome editing. Nature 516, 428–431 (2014).

30. Kohl, N.E. et al. Selective inhibition of ras-dependent transformation by a farnesyltransferase inhibitor. Science 260, 1934–1937 (1993).

31. Rowinsky, E.K., Windle, J.J. & Von Hoff, D.D. Ras protein farnesyltransferase: a strategic target for anticancer therapeutic development. Journal of Clinical Oncology 17, 3631–3652 (1999).

32. Collisson, E. et al. Comprehensive molecular profiling of lung adenocarcinoma: The cancer genome atlas research network. Nature 511, 543–550 (2014).

33. Feldser, D.M. et al. Stage-specific sensitivity to p53 restoration during lung cancer progression. Nature 468, 572–575 (2010).

34. Murray, C.W. et al. An LKB1–SIK axis suppresses lung tumor growth and controls differentiation. Cancer discovery 9, 1590–1605 (2019).

35. Tsherniak, A. et al. Defining a cancer dependency map. Cell 170, 564–576. e516 (2017).

36. Staffas, A., Karlsson, C., Persson, M., Palmqvist, L. & Bergo, M. Wild-type KRAS inhibits oncogenic KRAS-induced T-ALL in mice. Leukemia 29, 1032–1040 (2015).

37. Ambrogio, C. et al. KRAS dimerization impacts MEK inhibitor sensitivity and oncogenic activity of mutant KRAS. Cell 172, 857–868. e815 (2018).

38. Kong, G. et al. Loss of wild-type Kras promotes activation of all Ras isoforms in oncogenic Kras-induced leukemogenesis. Leukemia 30, 1542–1551 (2016).

39. Burgess, M.R. et al. KRAS allelic imbalance enhances fitness and modulates MAP kinase dependence in cancer. Cell 168, 817–829. e815 (2017).

40. Young, A., Lou, D. & McCormick, F. Oncogenic and wild-type Ras play divergent roles in the regulation of mitogen-activated protein kinase signaling. Cancer discovery 3, 112–123 (2013).

41. Grabocka, E. et al. Wild-type H-and N-Ras promote mutant K-Ras-driven tumorigenesis by modulating the DNA damage response. Cancer cell 25, 243–256 (2014).

42. Zhou, Y. et al. Signal integration by lipid-mediated spatial cross talk between Ras nanoclusters. Molecular and cellular biology 34, 862–876 (2014).

43. Zhou, Y. & Hancock, J.F. Ras nanoclusters: Versatile lipid-based signaling platforms. Biochimica et Biophysica Acta (BBA)-Molecular Cell Research 1853, 841–849 (2015).

44. Henis, Y.I., Hancock, J.F. & Prior, I.A. Ras acylation, compartmentalization and signaling nanoclusters. Molecular membrane biology 26, 80–92 (2009).

45. Inouye, K., Mizutani, S., Koide, H. & Kaziro, Y. Formation of the Ras dimer is essential for Raf-1 activation. Journal of Biological Chemistry 275, 3737–3740 (2000).

46. Muratcioglu, S. et al. GTP-dependent K-Ras dimerization. Structure 23, 1325–1335 (2015).

47. Lin, W.-C. et al. H-Ras forms dimers on membrane surfaces via a protein–protein interface. Proceedings of the National Academy of Sciences 111, 2996–3001 (2014).

48. Güldenhaupt, J. et al. N-Ras forms dimers at POPC membranes. Biophysical journal 103, 1585–1593 (2012).

49. Nan, X. et al. Ras-GTP dimers activate the mitogen-activated protein kinase (MAPK) pathway. Proceedings of the National Academy of Sciences 112, 7996–8001 (2015).

50. Terrell, E.M. et al. Distinct binding preferences between ras and raf family members and the impact on oncogenic ras signaling. Molecular cell 76, 872–884. e875 (2019).

51. Dankort, D. et al. A new mouse model to explore the initiation, progression, and therapy of BRAFV600E-induced lung tumors. Genes & development 21, 379–384 (2007).

52. Dietrich, P. et al. Neuroblastoma RAS viral oncogene homolog (NRAS) is a novel prognostic marker and contributes to sorafenib resistance in hepatocellular carcinoma. Neoplasia 21, 257–268 (2019).

53. Weyandt, J.D. et al. Wild-type Hras suppresses the earliest stages of tumorigenesis in a genetically engineered mouse model of pancreatic cancer. PloS one 10, e0140253 (2015).

54. To, M.D., Rosario, R., Westcott, P.M., Banta, K.L. & Balmain, A. Interactions between wild-type and mutant Ras genes in lung and skin carcinogenesis. Oncogene 32, 4028–4033 (2013).

55. Weyandt, J.D., Carney, J.M., Pavlisko, E.N., Xu, M. & Counter, C.M. Isoform-Specific effects of wild-type ras genes on carcinogen-Induced lung tumorigenesis in mice. Plos one 11, e0167205 (2016).

56. Jeng, H.-H., Taylor, L.J. & Bar-Sagi, D. Sos-mediated cross-activation of wild-type Ras by oncogenic Ras is essential for tumorigenesis. Nature communications 3, 1–8 (2012).

57. Miller, M.S. & Miller, L.D. RAS mutations and oncogenesis: not all RAS mutations are created equally. Frontiers in genetics 2, 100 (2012).

58. Xu, Q. & Dunbrack, R.L. ProtCID: a data resource for structural information on protein interactions. Nature communications 11, 1–16 (2020).

59. Alford, R.F. et al. The Rosetta all-atom energy function for macromolecular modeling and design. Journal of chemical theory and computation 13, 3031–3048 (2017).

60. Shapovalov, M.V. & Dunbrack Jr, R.L. A smoothed backbone-dependent rotamer library for proteins derived from adaptive kernel density estimates and regressions. Structure 19, 844–858 (2011).

